# RUNX1 Isoforms Regulate RUNX1 and Target-Genes Differentially in Platelets-Megakaryocytes: Association with Clinical Cardiovascular Events

**DOI:** 10.1101/2024.06.18.599563

**Authors:** Liying Guan, Deepak Voora, Rachel Myers, Fabiola Del Carpio-Cano, A. Koneti Rao

**Author notes:** **Correspondence** A. Koneti Rao, M.B.B.S. Sol Sherry Thrombosis Research Center and Section of Hematology Lewis Katz School of Medicine at Temple University 3420 N. Broad St., 204 MRB Philadelphia, PA 19140, USA.

## Abstract

**Background:** Hematopoietic transcription factor RUNX1 is expressed from proximal P2 and distal P1 promoter to yield isoforms RUNX1 B and C, respectively. The roles of these isoforms in RUNX1 autoregulation and downstream-gene regulation in megakaryocytes and platelets are unknown.

**Objectives:** To understand the regulation of RUNX1 and its target genes by RUNX1 isoforms.

**Methods:** We performed studies on RUNX1 isoforms in megakaryocytic HEL cells and HeLa cells (lack endogenous RUNX1), in platelets from 85 healthy volunteers administered aspirin or ticagrelor, and on the association of RUNX1 target genes with acute events in 587 patients with cardiovascular disease (CVD).

**Results:** In chromatin immunoprecipitation and luciferase promoter assays, RUNX1 isoforms B and C bound and regulated P1 and P2 promoters. In HeLa cells RUNX1B decreased and RUNX1C increased P1 and P2 activities, respectively. In HEL cells, RUNX1B overexpression decreased RUNX1C and RUNX1A expression; RUNX1C increased RUNX1B and RUNX1A. RUNX1B and RUNX1C regulated target genes (*MYL9, F13A1, PCTP, PDE5A* and others) differentially in HEL cells. In platelets RUNX1B transcripts (by RNAseq) correlated negatively with RUNX1C and RUNX1A; RUNX1C correlated positively with RUNX1A. RUNX1B correlated positively with *F13A1, PCTP, PDE5A, RAB1B*, and others, and negatively with *MYL9*. In our previous studies, RUNX1C transcripts in whole blood were protective against acute events in CVD patients. We found that higher expression of RUNX1 targets *F13A1* and *RAB31* associated with acute events.

**Conclusions:** RUNX1 isoforms B and C autoregulate RUNX1 and regulate downstream genes in a differential manner and this associates with acute events in CVD.

**Scientific category:** Platelets

**Essentials:**

1. RUNX1 is expressed from 2 promoters (P1 and P2) to yield isoforms RUNX1C and RUNX1B.
2. RUNX1B and RUNX1C regulate RUNX1 and target genes differentially in megakaryocytes/platelets.
3. In platelets RUNX1B and RUNX1C expression is inversely related and ticagrelor increases RUNX1C
4. RUNX1 target gene (*F13A1, RAB31*) expression in blood is associated with death or MI in cardiac disease.

## INTRODUCTION

Runt-related transcription factor 1 (RUNX1) is a master regulator of hematopoiesis, megakaryocyte (MK) and platelet biology [1–5]. Germline *RUNX1* haplodeficiency is associated with thrombocytopenia, platelet dysfunction and a striking predisposition to myeloid malignancies, referred to as familial platelet disorder with predisposition to myeloid malignancy (FPDMM) [6–10]. The *RUNX1* gene, located on chromosome 21, is regulated by two distinct promoters which are ∼160k base pairs (bp) apart: a distal P1 and a proximal P2 promoter [2, 11]. There are multiple RUNX1 isoforms, of which the best recognized are RUNX1B, RUNX1C, and RUNX1A [2, 12]. RUNX1A and RUNX1B are expressed from the P2 promoter while RUNX1C is from the P1 promoter [7, 13–16]. All three isoforms contain a DNA-binding RUNT domain; RUNX1A lacks the C-terminal transactivation domain [2, 13]. In Homo sapiens RUNX1B and RUNX1C differ by 32 N-terminus amino acids (AA), which regulate post-translational modifications and interaction with target DNAs and transcriptional co-regulators [17]. RUNX1 expression from 2 distinct promoters results in different 5’ and 3’ untranslated regions, which affects mRNA stability, translation activity and post-transcriptional modification [13, 17]. RUNX1 can both activate and repress target genes [17, 18].

RUNX1 is critical for hematopoiesis during embryogenesis and is required for the endothelial-to-hematopoietic transition [2, 19]. The differential expression patterns and role of RUNX1 isoforms have been explored. In mouse embryonic stem cell lines, RUNX1B is predominantly required for the initiation and ongoing hematopoietic development, while RUNX1C expression occurs later [20, 21]. In human studies, the roles of RUNX1 isoforms are unclear. While one study [22] found a similar sequential expression of RUNX1 isoforms and crucial importance of RUNX1B in hematopoietic development, another [23] showed that RUNX1C is the only isoform that is upregulated in hemogenic endothelium. During adult hematopoiesis, RUNX1C is predominantly expressed, while RUNX1B upregulation determines megakaryocytic and erythroid lineage differentiation [14]. RUNX1C has been shown to be important in MK progenitor proliferation and survival [24]. RUNX1A has also been shown to play an important role. RUNX1A enhances hematopoietic lineage commitment from human stem cells [25]. It is overexpressed in CD34+ cells in myeloproliferative neoplasms [26] and an altered RUNX1A/RUNX1C ratio has been linked to increased leukemia risk in trisomy 21 [27]. These studies point to differential functions of RUNX1 isoforms.

RUNX1 is reported to autoregulate its expression by binding to the P1 promoter in HL-60 cells and Jurkat T-cells [15]. However, the RUNX1 isoform involved in this autoexpression was not specified and the P2 promoter was not studied. Available evidence suggests that RUNX1 isoforms regulate the target genes differentially [28, 29]. Our studies using gene arrays showed that transcripts for RUNX1 target gene *PCTP* (phosphatidylcholine transfer protein) in whole blood correlate negatively with RUNX1 expression from P1 but not the P2 promoter [28]. Little is known regarding the effects of RUNX1 isoforms on its autoregulation and on target gene expression in MK/platelets.

Our goals in the present studies were to investigate the differential effects of RUNX1 isoforms on RUNX1 autoregulation and target gene regulation in MK/platelets. Expression profiling of platelets of patients with germline RUNX1 haplodeficiency (RHD) using Affymetrix microarrays have shown that numerous genes were downregulated or upregulated [30, 31]. Here, we focus on RUNX1 target genes, including *PCTP* [28]*, MYL9* (myosin light chain 9) [32]*, F13A1* (coagulation factor XIII A subunit) [33], *PDE5A* (phosphodiesterase 5A) (Guan and Rao, Unpublished), *RAB1B* ([34] and *RAB31* [35]. We provide evidence that the isoforms differentially autoregulate RUNX1 expression and regulate target genes in an isoform-specific manner. We show in human platelets that there is an inverse relationship between the expression of RUNX1B and RUNX1C, and that RUNX1B transcripts correlate with target gene expression, providing *in vivo* evidence for isoform-specific effects. In healthy volunteers, we show that a four-week daily exposure to platelet P2Y12 receptor antagonist ticagrelor modulates platelet RUNX1 expression, with an increase in RUNX1C transcripts but not RUNX1B, and expression of target genes. In patients with CVD, we have shown [29] that higher transcript levels from the RUNX1 P1 promoter had a protective effect with respect to death and MI. We now show that platelet expression of target genes *F13A1* and *RAB31* correlates with acute CV events. Overall, RUNX1 isoforms have differential effects and these studies highlight the importance of assessing isoform-specific effects in clinical scenarios where RUNX1 is involved and in its pharmacologic modulation for therapeutic purposes in FPDMM [36–38].

## METHODS

### Cells

Studies were performed in human erythroleukemia (HEL) cells and human cervical carcinoma (HeLa) cells, which do not express RUNX1 [15]. Details are provided in the Supplemental information.

### Chromatin immunoprecipitation and luciferase reporter studies

These were performed in HEL and HeLa cells as described in the Supplemental methods.

### Effect of RUNX1 isoforms on RUNX1 autoregulation and target genes

To study the effect of RUNX1B or RUNX1C overexpression on RUNX1 autoregulation and downstream gene expression, PMA-treated HEL cells (6x10^5^) or HeLa cells (6x10^5^) cultured in 6-well plate were transfected with different concentrations of RUNX1B or RUNX1C expression vector. After 48 hours, cell lysates were subjected to immunoblotting using antibodies against RUNX1, RUNX1C, PCTP, MYL9, F13A1, PDE5A and GAPDH. Secondary antibodies were IRDye-labeled antibodies (LI-Cor Biosciences). Immunoreactive proteins were detected with the Odyssey Infrared Imaging System (LI-Cor Biosciences). The density of the bands was measured using Odyssey V3.0 (LI-Cor Biosciences). Protein levels were normalized to GAPDH. RNA extracted using the RNeasy Mini Kit was used for cDNA synthesis using SuperScript IV First Strand Synthesis System. The expression of RUNX1 isoforms and downstream genes was analyzed by qPCR using PowerUp SYBR Green Master Mix (primers listed in Supplemental Table 1). Reactions were performed using the Mastercycler ep realplex (Eppendorf) system, and Ct values for each gene expression were generated. Gene expression was determined using the ΔΔCt method with GAPDH as reference gene.

### Studies in human subjects on platelet RUNX1 isoforms and target genes, and on the effects of ticagrelor and aspirin

Samples were obtained from a previously reported [39, 40] prospective, open label, randomized cross-over study to assess the effects of antiplatelet therapy (non-enteric coated aspirin 81 mg/day, aspirin 325 mg/day, and ticagrelor 90 mg twice daily) on platelet function and gene expression. Briefly, 85 healthy volunteers were enrolled for a baseline platelet RNA collection and platelet function assessments (V1) and randomized to either 4 weeks of 81 mg/day vs. 325 mg/day aspirin after which they had repeated baseline assessments (V2) and crossed over to the other aspirin dose for an additional 4 weeks (V3). After a 4-week washout (V4), participants were loaded with 180 mg ticagrelor and prescribed 90 mg twice daily ticagrelor and returned for their final assessments (V5). Platelets were isolated from whole blood using a leukocyte depletion procedure [41]. Purified platelet RNA was collected before and after each exposure and analyzed by RNA sequencing [40]. The Duke Institutional Review Board approved the study protocol.

### Platelet RNA quantification and statistical analyses

Sequence data processing and alignment, and differential expression by aspirin and ticagrelor treatment are described in detail previously [39, 40] and in the Supplement.

### Association of RUNX1 isoform-specific expression with RUNX1 target genes

Expression of RUNX1 isoforms and the RUNX1 target genes was quantified as log2 counts per million reads mapped (log2 CPM). Two association models were tested: 1. Model A (off treatment): Target gene ∼ RUNX1 isoform + (1|*subject*), and 2. Model B (on and off treatment): Target gene ∼ Runx1 isoform + treatment + (1|*subject*). Both models were estimated using linear mixed-effect regression with a random intercept for subject to account for repeated sampling within a person. The Benjamini-Hochberg method was used to adjust p-values for multiple testing across the RUNX1 isoforms and target genes, within each model. Model A was repeated genome-wide (14,333 unique transcripts) and the resulting association test statistics (absolute value of the t-statistics) were used to assess for enrichment of RUNX1-gene correlation within gene sets that are downregulated (228 genes) and upregulated (21 genes) in platelets from our RHD patient [30] using a two-sample t-test. All platelet RNA analyses were conducted in the statistical program R using the packages limma and edgeR, or Ime4.

### Studies on patients with cardiovascular disease

Methods used for studies in CVD patients referred for cardiac catheterization and followed for CV events, and their molecular data have been previously described [42] and provided in the Supplemental Information. Briefly, combined data from two cohorts of CVD patients (one case-control (n=190) and another observational (n=397)), baseline clinical, medication, and peripheral blood microarray data, and long-term clinical outcomes from the Duke Catheterization Genetics (CATHGEN) cohort were used for these analyses.

Statistical analyses of peripheral blood gene expression data have been previously described [28]. Briefly, to assess for the association between target gene *(F13A1*, *RAB31*, *PDE5A*, or *RAB1B*) expression and CV events, we used two independent longitudinal follow up cohorts from the CATHGEN study (case-control and observational). In each cohort, each probe set was tested for association with the combined outcome of death or MI events using linear regression, controlling for the effects of age, sex, and race on death or MI outcomes. All CATHGEN data processing and statistical analyses were conducted in R version 3.2 (http://www.r-project.org/) using the packages affy, meta, and limma for normalization, meta-analyses, and moderated t-tests, respectively. Additional details regarding the R programming required to process and analyze the RNA-seq data is described in the GEO submission of these data: GSE158765.

### Statistical analysis for other studies

Results were expressed as mean ± SEM. Differences were compared using Student’s t-test or one- and two-way ANOVA, using the GraphPad Prism, version 8 (GraphPad Software) and considered significant at *P* < 0.05.

## RESULTS

### RUNX1B and RUNX1C bind to P1 and P2 promoters and regulate activity

RUNX1 is expressed from two promoters [1, 2, 11, 12]; RUNX1C is expressed from P1 while RUNX1B and RUNX1A are from the P2 promoter (Figure 1A). RUNX1C has 32 additional AA at the N-terminus compared to RUNX1B (Figure 1B). We developed a rabbit antibody that binds to the 16 AA N-terminus of RUNX1C, referred to hereafter as the RUNX1C-specific antibody. Studies were done with the commercially available antibodies (referred to as RUNX1 antibodies), which bind to all three isoforms and with the RUNX1C-specific antibody. In HeLa cells, which lack endogenous RUNX1 [15], none of the antibodies detected RUNX1 (Figure 1C), while in HEL cells the RUNX1 antibodies detected bands corresponding to RUNX1 B (48 kDa) and RUNX1C (52 kDa). Upon expressing RUNX1B or RUNX1C in HeLa cells, the RUNX1 antibodies detected only the appropriate isoform, and the RUNX1C-specific antibody detected the band corresponding to RUNX1C but not RUNX1B (Figure 1C). This validated the specificity of the RUNX1C-specific antibody.

**Figure 1.**
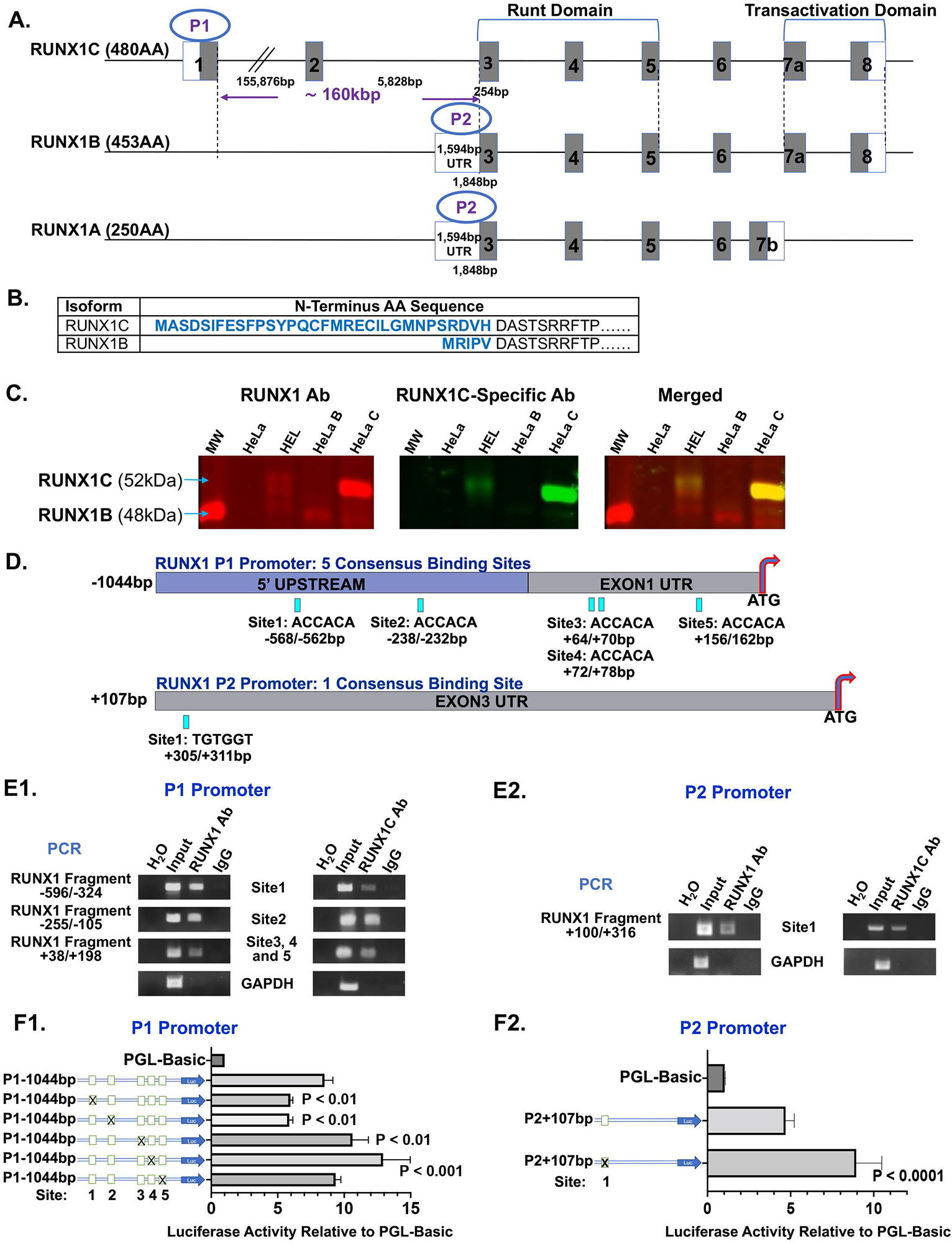
RUNX1 P1 and P2 promoter regions and autoregulation by RUNX1 isoforms. **A. Schematic representation of the three major isoforms of the human RUNX1 transcript.** The locations of promotor regions P1 and P2 are shown. Functional exons encoding the DNA-binding domain (Runt Domain) and transactivation domain are indicated. AA, amino acids; UTR, untranslated region. **B. Amino acid (AA) sequence of Homo sapiens RUNX1C and RUNX1B N-terminus.** RUNX1C and RUNX1B are transcribed from two distinct promoters. RUNX1C has additional 32 AA at the N-terminus compared to RUNX1B. The N-terminus AA sequence of RUNX1B and RUNX1C are shown. Those different between RUNX1C and RUNX1B are shown in bold blue. **C. Validation of the RUNX1C-specific antibody.** Shown are immunoblots for RUNX1B and RUNX1C in lysates of HeLa and HEL cells. HeLa cells have negligible RUNX1 protein; HEL cells express both RUNX1B and RUNX1C. We transiently expressed RUNX1B (HeLaB) and RUNX1C (HeLaC) separately in HeLa cells. Shown are immunoblots using commercial RUNX1 antibodies that bind to both RUNX1 isoforms (left panel, red) and RUNX1C-specific antibodies developed by us (middle panel, green). The right panel shows the merged image. MW, molecular weight; Ab, antibodies. **D. The 5’ upstream region of RUNX1 P1 (top) and P2 (bottom) promoter regions with the RUNX1 consensus binding sites.** There are five RUNX1 consensus binding sites in the P1 upstream promoter region and one RUNX1 consensus binding site in the P2 promoter region. UTR, untranslated region. **E. Binding of RUNX1 and RUNX1C to P1 (Panel E1) and P2 (Panel E2) promoter regions by chromatin immunoprecipitation (ChIP).** HEL cells were differentiated with PMA (30 nM) for 48 hours. ChIP was performed using anti-RUNX1 antibody (RUNX1 Ab), house generated anti-RUNX1C antibody (RUNX1C Ab) or control IgG. PCR was performed using primers shown in Supplemental Table 1 to amplify RUNX1 promoter regions encompassing RUNX1 binding sites and GAPDH. Each panel shows PCR products obtained from amplification of water (H_2_O), total input DNA, anti-RUNX1 antibody- or anti-RUNX1C antibody-immunoprecipitated samples, and IgG-immunoprecipitated samples. **F. Luciferase reporter studies on the RUNX1 P1 (Panel F1) and P2 (Panel F2) promoters in PMA-treated HEL cells.** P1 promoter region (1.1 kbp) has five RUNX1 consensus binding sites (F1) and P2 promoter region (1.5kbp) has one RUNX1 binding site (F2). Mutant promoters of each binding site were generated and indicated with “X” at the site. HEL cells were transfected with either wild-type or mutated RUNX1 promoter luciferase reporter vectors. Cells were transfected with promoterless PGL-Basic plasmid as a baseline control (set as 1 for standardization). Cells were harvested 24 hours later and cell lysates were evaluated for luciferase activity using the dual luciferase reporter system. Data were normalized with Renilla luciferase values used as an internal control. Statistical analysis was performed using one-way ANOVA (n=6, mean±SEM). *P* values indicate comparisons of the activity of each mutant construct with that of the wild-type promoter luciferase construct.

All three RUNX1 isoforms share a homologous Runt Domain (Figure 1A) that mediates the binding of target DNAs [15]. *In silico* analysis revealed five RUNX1 consensus binding sites within 600 bp of the P1 5’ upstream region from exon1 and one RUNX1 site in the UTR of exon3 (Figure 1D). We performed ChIP assay using PMA-treated HEL cells and antibodies recognizing both RUNX1B and RUNX1C, and separately with RUNX1C-specific antibodies. Both sets of antibodies but not the control IgG enriched by PCR amplification the regions encompassing the RUNX1 consensus sites in the P1 and P2 promoters (Figure 1E). In the P1 promoter, RUNX1 bound to regions encompassing sites 1 and 2, and the region with sites 3, 4 and 5, which are in close proximity (Figure 1E1). In the P2 promoter, both the RUNX1 antibodies and the RUNX1C-specific antibodies bound to the region with the single consensus site (Figure 1E2). Thus, RUNX1 binds to both promoters, particularly RUNX1C. Mutations of sites 1 and 2 in the P1 promoter decreased promoter activity (Figure 1F1); while mutations of sites 3, 4 increased activity. In the P2 promoter, mutation of the single binding site increased activity (Figure 1F2). These studies indicate that RUNX1 binds to the P1 and P2 promoters to regulate transcriptional activities.

### Studies in HeLa cells: RUNX1B increases and RUNX1C decreases P1 and P2 promoter activities

We examined isoform-specific regulation of promoter activities by co-transfecting each isoform along with the P1 or P2 promoter luciferase vector in HeLa cells, which lack endogenous RUNX1. We co-transfected the P1-promoter luciferase vector with increasing amounts of RUNX1B or RUNX1C expression vector (Figure 2A). RUNX1B induced a dose-dependent decrease in P1 activity; RUNX1C showed a dose-dependent increase (Figure 2A). In MK, RUNX1B and RUNX1C are co-expressed. To assess the interaction between RUNX1B and RUNX1C on P1 promoter activity, we expressed a constant amount of RUNX1C and an increasing amount of RUNX1B expression vector along with the P1-promoter. RUNX1B decreased P1 promoter activity induced by RUNX1C in a dose dependent manner (Figure 2A, right panel). Thus, RUNX1B and RUNX1C compete in regulating P1 promoter activity. The same studies were conducted with the P2 promoter (Figure 2B). RUNX1B decreased P2 activity in a dose-dependent manner. With RUNX1C, we did not observe a dose-dependent alteration in P2 activity (Figure 2B). However, with constant RUNX1B expression, which decreased activity by itself, added RUNX1C increased P2 activity in a dose-dependent manner (Figure 2B, right panel). This suggests that even though we were unable to show a direct effect on the P2 promoter, RUNX1C interacts with this promoter to compete with RUNX1B.

**Figure 2.**
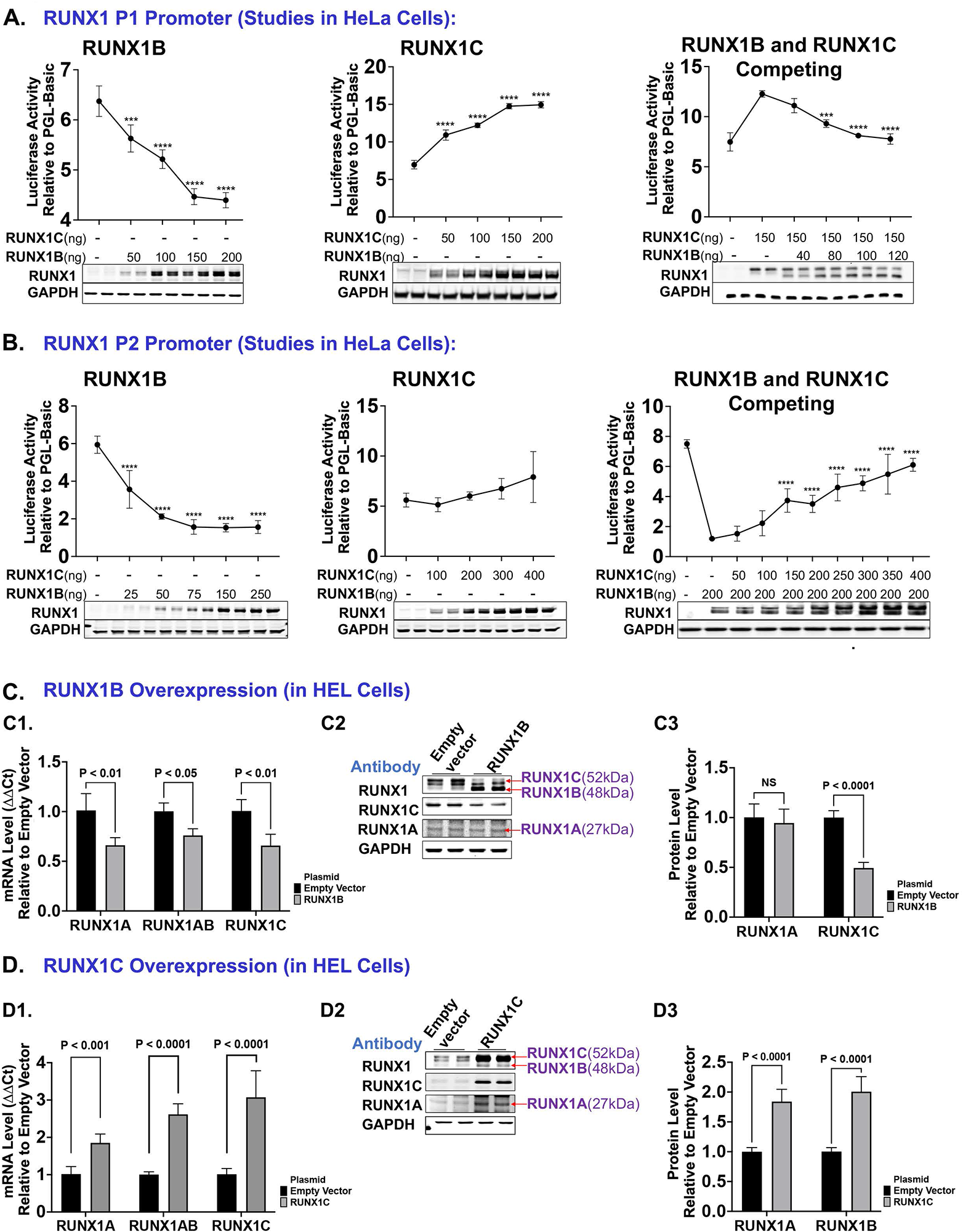
Differential effects of RUNX1 isoforms B and C on RUNX1 expression in HeLa cells and HEL cells. **A. Effect of RUNX1B and RUNX1C on P1 promoter activity in HeLa Cells.** HeLa cells were transiently co-transfected with 800 ng of RUNX1 P1 promoter luciferase reporter vector and indicated amounts of RUNX1B (left), RUNX1C (middle) or both expression vectors (right). Cells were transfected with 800 ng promoterless PGL-Basic plasmid as a baseline control (set as 1 for standardization). They were harvested 24 hours later for luciferase reporter assay and immunoblotting to confirm RUNX1 transfection levels. Luciferase activity was normalized with Renilla luciferase as an internal control. Statistical analysis was performed using one-way ANOVA (n=6, mean±SEM, ***p<0.001, ****p<0.0001). RUNX1 protein was detected by probing blots with anti-RUNX1 antibody. GAPDH is shown as an internal loading control. **B. Effect of RUNX1B and RUNX1C on P2 promoter activity in HeLa Cells.** HeLa cells were transiently co-transfected with 800 ng RUNX1 P2 promoter luciferase reporter vector and indicated amounts of RUNX1B (left), RUNX1C (middle) or both expression vectors (right). Cells were transfected with 800 ng promoterless PGL-Basic plasmid as a baseline control (set as 1 for standardization). Cells were harvested 24 hours later for luciferase reporter assay and immunoblotting to confirm RUNX1 transfection levels. Luciferase activity was normalized with Renilla luciferase as an internal control. Statistical analysis was performed using one-way ANOVA (n=6, mean±SEM, ****p<0.0001). RUNX1 protein was detected by probing blots with anti-RUNX1 antibody. GAPDH is shown as an internal loading control. **C. Effect of RUNX1B overexpression on RUNX1 isoforms in PMA-treated HEL cells.** HEL cells treated with 30 nM PMA were transfected with 900 ng pCMV6-RUNX1B expression vector or pCMV6 empty vector. C1, cells were extracted for total mRNAs and subjected to RT-qPCR. Shown are relative mRNA expression levels for endogenous RUNX1 isoforms using the ΔΔCt method. Results from cells transfected with the empty vector are set as 1. GAPDH is used as an internal control. C2, cells were extracted for immunoblotting to assess RUNX1 isoform protein levels; shown are immunoblots of cell lysates for RUNX1 (using commercial RUNX1 antibody), RUNX1C (using RUNX1C-specific antibody), RUNX1A (using commercial RUNX1 antibody) and GAPDH as a loading control. C3, protein band intensities were measured using Odyssey V3.0 and normalized to GAPDH. The intensities from samples transfected with the empty vector were set as 1 for normalization. Statistical analysis was performed using Student’s t-test (n=6, mean±SEM). **D. Effect of RUNX1C overexpression on RUNX1 isoforms in PMA-treated HEL cells.** HEL cells were transfected with 900 ng M02-RUNX1C expression vector or M02 empty vector. D1, cells were extracted for total mRNAs and subjected to RT-qPCR. Shown are relative mRNA expression levels of endogenous RUNX1 isoforms using the ΔΔCt method. Results from cells transfected with the empty vector are set as 1. GAPDH is used as an internal control. D2, RUNX1 isoform protein levels; shown are immunoblots of cell lysates for RUNX1 (using commercial RUNX1 antibody), RUNX1C (using RUNX1C specific antibody), RUNX1A (using commercial RUNX1 antibody) with GAPDH as a loading control. D3, protein band intensities were measured using Odyssey V3.0 and normalized to GAPDH; the intensities from samples transfected with the empty vector were set as 1 for standardization. Statistical analysis was performed using Student’s t-test (n=6, mean±SEM).

### Studies in megakaryocytic HEL cells: RUNX1B increases and RUNX1C decreases P1 and P2 promoter expression

In addition to RUNX1 isoforms B and C, MK and HEL cells have RUNX1A, which is expressed from the P2 promoter [12, 21]. We studied the effects of overexpressing RUNX1B (Figure 2C) and RUNX1C (Figure 2D) in HEL cells by assessing mRNA and protein levels of the isoforms. Because RUNX1B transcripts share a homologous N-terminus with RUNX1A and share a homologous C-terminus with RUNX1C (Figure 1A), we were unable to measure individual RUNX1B transcript levels using traditional qPCR. Therefore, we examined combined RUNX1A and RUNX1B (RUNX1AB) mRNA levels. RUNX1B overexpression decreased RUNX1A, RUNX1AB and RUNX1C mRNA levels compared to control empty vector (Figure 2C1). RUNX1C protein was decreased ∼50% by immunoblotting using RUNX1 antibodies and the RUNX1C-specific antibodies; RUNX1A protein was unchanged (Figure 2C2 and 2C3). Because ectopically expressed RUNX1B overlaps with endogenous RUNX1B on immunoblotting (Figure 2C2), the endogenous RUNX1B expression was indirectly assessed using transcript levels for RUNX1AB (P2 promoter expression) and RUNX1A. Both transcripts were decreased, suggesting inhibition of RUNX1B expression as well, because RUNX1B transcripts constitute the bulk of RUNX1AB transcripts (data not shown). Thus, RUNX1B overexpression inhibits expression of all three isoforms.

RUNX1C overexpression increased mRNA for RUNX1A ∼1.5-fold; RUNX1AB ∼2.5-fold and RUNX1C 3-fold (Figure 2D1). The increase in RUNX1AB suggested that endogenous RUNX1B mRNA was also increased; on immunoblotting, there was a 2-fold increase in both RUNX1A and RUNX1B (Figure 2D2 and 2D3). These studies indicate that the isoforms regulate their own expression differentially in megakaryocytic cells with RUNX1B inhibiting, and RUNX1C enhancing expression from both promoters.

### RUNX1 isoforms differentially regulate target gene expression in HeLa and HEL cells

RUNX1 regulates numerous genes [3, 43]. In HeLa cells, RUNX1B overexpression increased *MYL9, PCTP* and *PDE5A* mRNA and protein in a dose-dependent manner (Figure 3A). RUNX1C overexpression decreased *MYL9* and *PCTP* mRNA and protein (Figure 3B) but not *PDE5A*. HeLa cells express negligible *F13A1* precluding studies on this gene.

**Figure 3.**
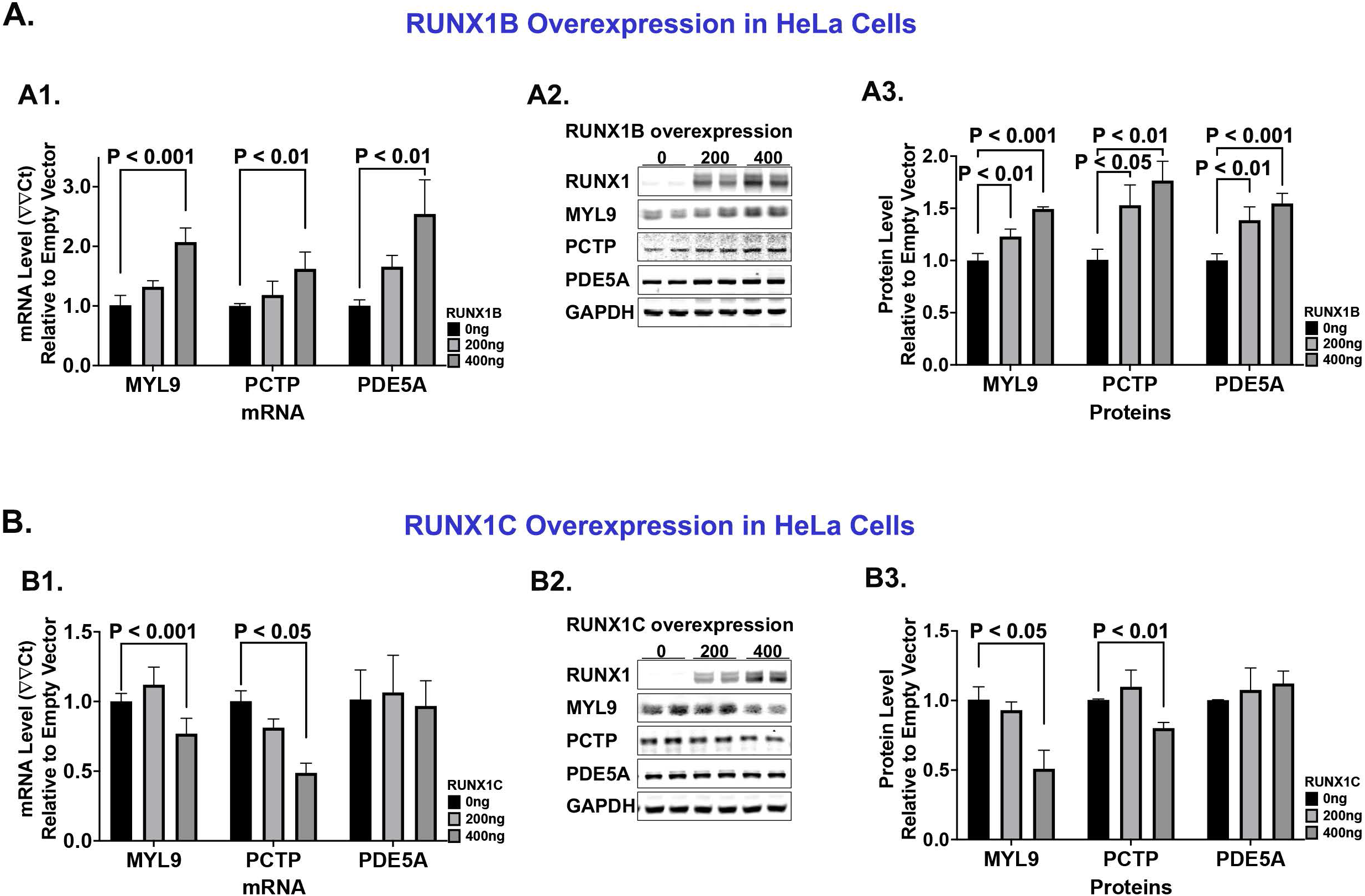
Effect of overexpression of RUNX1B and RUNX1C on target genes in HeLa cells. **A. Effect of RUNX1B overexpression in HeLa cells.** HeLa cells were transfected with the pCMV6-RUNX1B expression vector or pCMV6 empty vector. A1, cells were extracted for total mRNAs and subjected to RT-qPCR. Shown are relative mRNA levels of downstream genes *MYL9, PCTP* and *PDE5A* using the ΔΔCt method. Results from cells transfected with the empty vector are set as 1. GAPDH was used as an internal control (n=4, mean±SEM). A2, shown are immunoblots of cell lysates for RUNX1, MYL9, PCTP, and PDE5A with GAPDH as a loading control. A3, protein band intensities were measured using Odyssey V3.0 and normalized to GAPDH; the intensities from samples transfected with the empty vector were set as 1 for standardization. Statistical analysis was performed using one-way ANOVA (n=4, mean±SEM). **B. Effect of RUNX1C overexpression in HeLa cells.** HeLa cells were transfected with the M02-RUNX1C expression vector or M02 empty vector. B1, cells were extracted for total mRNAs and subjected to RT-qPCR; shown are relative mRNA levels of *MYL9, PCTP* and *PDE5A* using the ΔΔCt method. Results from cells transfected with the empty vector are set as 1. GAPDH was used as an internal control (n=4, mean±SEM). B2, shown are immunoblots of cell lysates for RUNX1, MYL9, PCTP and PDE5A with GAPDH as a loading control. B3, quantification of protein band intensities normalized to GAPDH; the intensities from samples transfected with the empty vector were set as 1 for standardization. Statistical analysis was performed using one-way ANOVA (n=4, mean±SEM).

In HEL cells, RUNX1B overexpression increased *F13A1, MYL9*, *PCTP* and *PDE5A* mRNA and protein levels (Figure 4A). RUNX1C overexpression decreased *MYL9* and *PCTP* mRNA and protein; *F13A1* or *PDE5A* were unchanged (Figure 4B). Thus, RUNX1B and RUNX1C differentially regulate downstream genes in HeLa cells and HEL cells.

**Figure 4.**
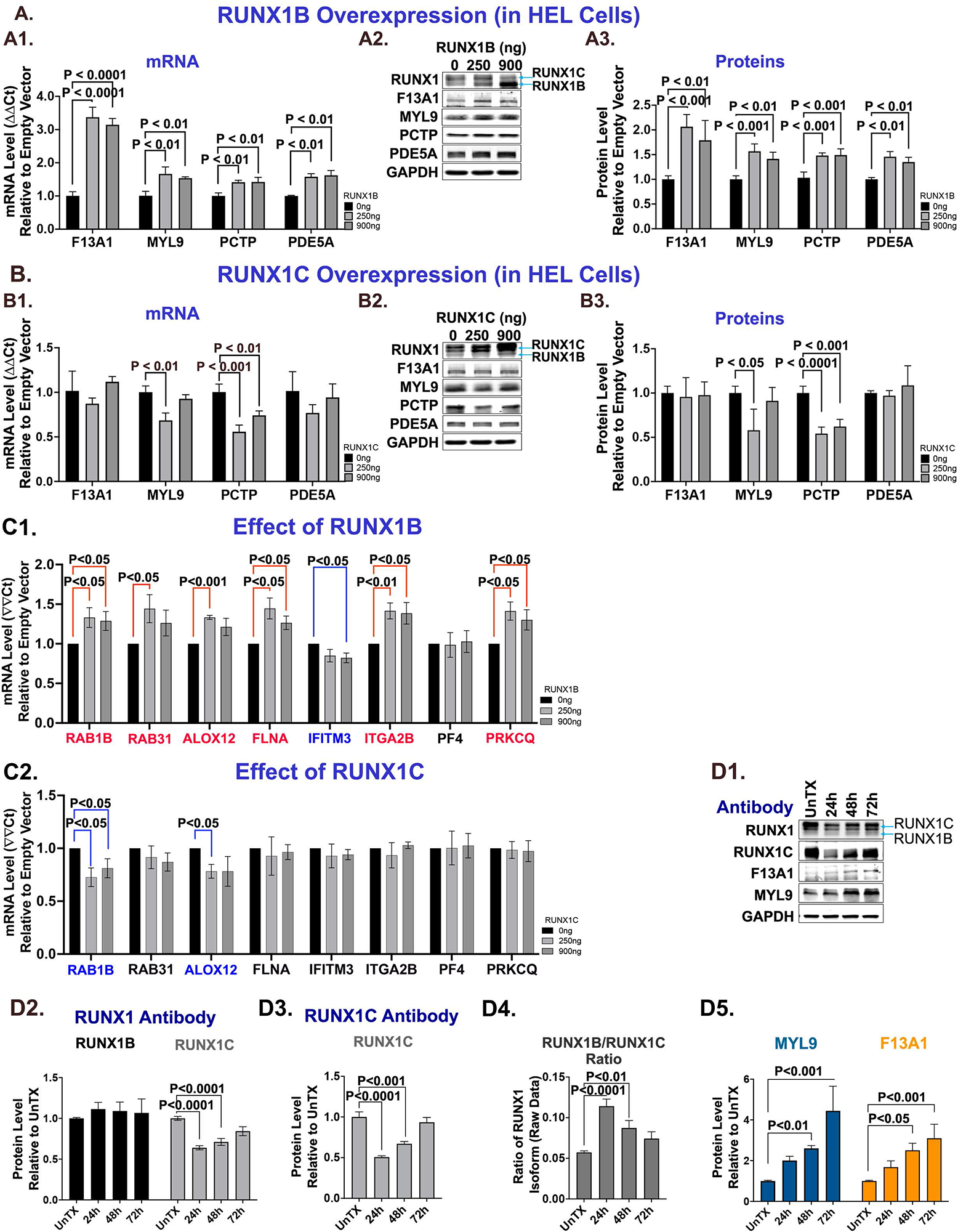
Effect of overexpression of RUNX1B and RUNX1C on target genes in HEL cells. **A. Effect of RUNX1B overexpression on target genes.** PMA-treated HEL cells were transfected with the pCMV6-RUNX1B expression vector or pCMV6 empty vector. A1, cells were extracted for total mRNA and subjected to RT-qPCR. Shown are relative mRNA levels of *F13A1, MYL9, PCTP* and *PDE5A* using the ΔΔCt method. Results from cells transfected with the empty vector are set as 1. GAPDH was used as an internal control (n=4, mean±SEM). A2, shown are immunoblots of cell lysates for RUNX1, F13A1, MYL9, PCTP and PDE5A with GAPDH as a loading control; A3, quantification of protein band intensities normalized to GAPDH; the intensities from samples transfected with the empty vector were set as 1 for standardization. Statistical analysis was performed using one-way ANOVA (n=4, mean±SEM,). **B. Effect of RUNX1C overexpression on target genes.** HEL cells were transfected with the M02-RUNX1C expression vector or M02 empty vector. B1, cells were extracted for total mRNA and subjected to RT-qPCR. Shown are relative mRNA levels of *F13A1, MYL9, PCTP* and *PDE5A* using the ΔΔCt method. GAPDH is used as an internal control (n=4, mean±SEM). B2, shown are immunoblots of cell lysates for RUNX1, F13A1, MYL9, PCTP, PDE5A and GAPDH as a loading control. B3, quantification of protein band intensities normalized to GAPDH; the intensities from samples transfected with the empty vector were set as 1 for standardization. Statistical analysis was performed using ANOVA (n=4, mean±SEM,). **C. Effect of RUNX1 isoforms on RUNX1 target gene expression in HEL cells using qPCR.** C1, HEL cells were transfected with the pCMV6-RUNX1B expression vector or pCMV6 empty vector; C2, HEL cells were transfected with the M02-RUNX1C expression vector or M02 empty vector. Cells were extracted for total mRNA and subjected to RT-qPCR. Shown are relative mRNA levels of *RAB1B, RAB31, ALOX12, FLNA, IFITM3, ITGA2B, PF4 and PRKCQ* using the ΔΔCt method. Results from cells transfected with the empty vectors are set as 1. GAPDH was used as an internal control. Upregulated genes are shown in red, and downregulated genes in blue. Statistical analysis was performed using one-way ANOVA (n=4, mean±SEM). **D. Effect of PMA-treatment of HEL cells on RUNX1 isoforms and target genes *MYL9* and *F13A1*.** HEL cells were treated with 30 nM PMA to induce megakaryocytic differentiation for up to 72 hours. Lysates were prepared from untreated HEL cells (UnTX), and at 24, 48 and 72 hours. D1, shown are immunoblots of cell lysates for RUNX1 (using commercial RUNX1 antibody), RUNX1C (using RUNX1C-specific antibody), F13A1, MYL9 and GAPDH. D2, protein band intensities were measured using Odyssey 3.0 and the intensities from untreated HEL cells (UnTX) were set as 1 for standardization. Bar graphs show protein levels of RUNX1B and RUNX1C relative to GAPDH using commercial RUNX1 antibody. D3, RUNX1C protein levels using RUNX1C-specific antibody, GAPDH was used as an internal control. Statistical analysis was performed using one-way ANOVA (n=3, mean±SEM). D4, ratio of RUNX1B to RUNX1C protein levels on PMA treatment. Protein band intensities were quantified. Shown are ratios of RUNX1B over RUNX1C protein levels using commercial RUNX1 antibody. Statistical analysis was performed using ANOVA (n=3, mean±SEM). D5, MYL9 and F13A1 protein levels on PMA treatment. Protein band intensities of MYL9 and F13A1 were quantified. Results from UnTX cells were set as 1 for standardization. Bar graphs show relative MYL9 and F13A1 protein levels using GAPDH as an internal control, Statistical analysis was performed using ANOVA, (n=3, mean±SEM).

To obtain further evidence, we examined the effects in HEL cells on eight additional genes (mRNA levels) whose platelet expression was altered in RHD patients [30, 31]. Seven were differentially regulated by RUNX1B and RUNX1C (Figure 4C). *RAB1B, RAB31, ALOX12, FLNA, ITGA2B, PRKCQ* were upregulated by RUNX1B (Figure 4C1); *RAB1B* and *ALOX12* were downregulated by RUNX1C, with the others unaffected (Figure 4C2). *IFITM3* was downregulated by RUNX1B and unaffected by RUNX1C. Neither altered *PF4*. Overall, 11 of 12 genes studied showed differential regulation by the isoforms.

Because PMA induces megakaryocytic differentiation of HEL cells [44] we examined its effects. With PMA treatment, RUNX1B protein levels appeared unchanged (Figure 4D1); RUNX1C decreased over 48-72 hours by immunoblotting using either the RUNX1 antibody (Figure 4D2) or the RUNX1C-specific antibody (Figure 4D3). The RUNX1B/RUNX1C ratio was increased (Figure 4D4) reflecting the relative RUNX1B increase. F13A1 and MYL9 proteins (Figure 4D5) increased over time, consistent with their upregulation by RUNX1B (Figure 4A).

### Studies in platelets: relationships between RUNX1 isoforms and target genes by RNAseq

To obtain *in vivo* evidence to support our findings in HEL cells (Figure 4), we assessed RUNX1 isoforms and 12 RUNX1 target genes [29] in leukocyte-poor platelets from 85 volunteers using RNAseq. Total RUNX1 transcripts were lower than those of target genes (Supplemental Figure 1A); RUNX1B and RUNX1C were similar, with RUNX1A being lower (Figure 5A). The ratios of RUNX1A/RUNX1C and RUNX1B/RUNX1C were similar (Supplemental Figure 1B). RUNX1B correlated negatively with RUNX1C and RUNX1A (Figure 5B). RUNX1C correlated positively with RUNX1A. These findings are in line with those in HEL and HeLa cells (Figure 2) and with the differential autoregulation by the isoforms.

**Figure 5.**
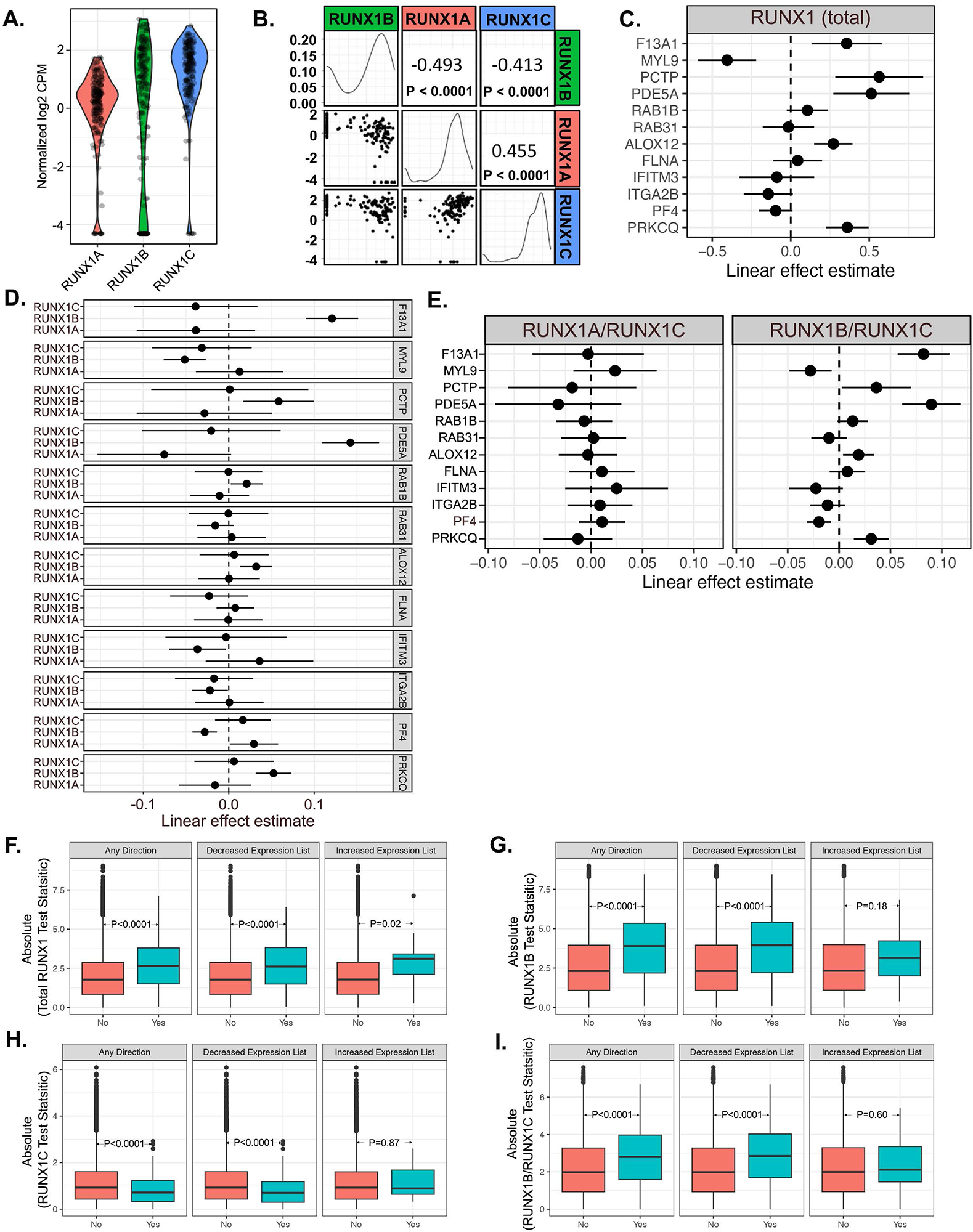
RUNX1 isoforms and their relationships to each other and target genes in platelets from healthy volunteers. **A. Relative expression of RUNX1 isoforms in platelets.** RNA was isolated from leukocyte-poor platelets from 85 healthy volunteers and analyzed using RNAseq as described in Methods. Shown are the Trimmed Mean of M component (TMM)-normalized expression represented as log2 counts per million (CPM) reads mapped of the three RUNX1 isoforms expression in baseline samples. **B. Relationships between RUNX1 isoforms in platelets.** The relationships between RUNX1 isoforms A, B and C are displayed. Scatterplots are shown for pairs of isoform TMM-normalized expression using log2 CPM values below the diagonal. The distribution of each isoform expression in log2 CPM is shown along the diagonal. Spearman rank correlation on log2 CPM expression values and p-values are displayed for each pair of isoforms. **C. Correlation between the expression of total RUNX1 and target genes in human platelets.** The correlation between total RUNX1 expression and its downstream genes was analyzed using linear mixed effects regression as described in the Methods section. Displayed are the linear effect estimates from the regression model and the 95% confidence interval, representing the increases in the target gene expression (*F13A1, MYL9, PCTP, PDE5A, RAB1B, RAB31, ALOX12, FLNA, IFITM3, ITGA2B, PF4*, and *PRKCQ*) for each unit increase in log2 normalized expression of total RUNX1. **D. Correlation between the expression of RUNX1 isoforms and target genes in platelets.** The correlation between each RUNX1 isoform and its downstream genes was estimated using linear mixed effects regression. Displayed are the linear effect estimates from the regression model and the 95% confidence interval, representing the increase in the target gene expression for each unit increases in log2 normalized expression of each RUNX1 isoform. **E. Correlation of ratios of RUNX1A to RUNX1C (left) and RUNX1B to RUNX1C (right) isoform with expression of downstream genes in platelets.** The correlation between the ratios (RUNX1A/RUNXC; RUNX1B/RUNX1C) and downstream genes was analyzed using linear mixed effects regression as described in the Methods section. Displayed are the linear effect estimates from the regression model and the 95% confidence interval, representing the increases in the target gene expression for each unit increase in log2 normalized expression of RUNX1A/RUNX1C and RUNX1B/RUNX1C. **F. Enrichment test results for total RUNX1 correlation within RHD regulated gene sets.** RNA was isolated from leukocyte-poor platelets from 85 healthy volunteers and analyzed for genome-wide gene expression using RNAseq. The resulting association test statistics (absolute value of the t-statistics) were used to assess the enrichment of total RUNX1 correlation within RHD regulated gene sets (228 decreased and 21 increased expression). Shown are the absolute RUNX1 gene test statistic values (y-axis) using a two-sample t-test across three categories: Any Direction, Decreased Expression List, and Increased Expression List. “No” indicates non-RHD regulated gene set. “Yes” indicates RHD regulated gene set. Enrichment p values generated from Fisher’s exact test are presented. **G-I. Enrichment test results for correlation of RUNX1B (G), RUNX1C (H), and RUNX1B/RUNX1C ratio (I) within RHD regulated gene sets.**

We examined the relationships of the isoforms to the 12 genes studied in HEL cells (Figure 4). In platelets, expression of 5 genes (*F13A1, PCTP, PDE5A, ALOX12, PRKCQ)* correlated positively with total RUNX1 transcripts (Figure 5C); there was an inverse correlation with *MYL9*. Six (*F13A1, PCTP, PDE5A, RAB1B, ALOX12, PRKCQ)* correlated positively with RUNX1B (Figure 5D), as noted in HEL cells (Figure 4). *IFITM3* was negatively regulated by RUNX1B in both HEL cells and platelets. Three (*MYL9, ITGA2B, PF4)* correlated negatively with RUNX1B (Figure 5D); two (*MYL9, ITGA2B)* were positively regulated in HEL cells (Figure 4). None correlated with RUNX1C or RUNX1A. We assessed the relationships between the RUNX1B/RUNX1C ratio and the 12 target genes (Figure 5E). They generally paralleled those between RUNX1B and the target genes (Figure 5D). The RUNX1A/RUNX1C ratio did not correlate with any of the genes (Figure 5E). Thus, in platelets RUNX1B expression correlates positively with most genes that were positively regulated by RUNX1B in HEL cells.

We have previously reported genes that are downregulated and upregulated in platelets from in our patient with RHD in studies using Affymetrix arrays [30]. Using genome-wide platelet RNAseq data from healthy volunteers, we correlated the expression of all expressed transcripts and the RUNX1 isoforms, RUNX1B/RUNX1C ratio and RUNX1A/RUNX1C ratio. Using these linear associations, we performed enrichment analysis for the downregulated, upregulated and combined gene sets identified in our RHD patient. There was significant enrichment for total RUNX1, RUNX1B, RUNX1C, RUNX1B/RUNX1C ratio (Figure 5 F-I) and RUNX1A/RUNX1C ratio (Supplemental Figure 2B) with the set of genes downregulated in the patient, but not with upregulated gene set, which has fewer genes (Supplemental Table 2-4). There was no enrichment for RUNX1A by itself (Supplemental Figure 2A). These enrichment analyses corroborate findings for RUNX1B using an expanded gene set. By using this expanded gene set, these data also suggest a role for RUNX1C that was not apparent with the selected target genes in HEL cell experiments.

### Effect of aspirin and ticagrelor on platelet RUNX1 and target gene expression by RNAseq

Our prior studies in HEL cells and in whole blood showed that RUNX1 is an aspirin-responsive gene. In HEL cells and in whole blood of healthy individuals administered aspirin for 4-weeks, it enhanced RUNX1 expressed from the P1 promoter and inhibited those from the P2 promoter [29]. Moreover, *in vitro* P2Y12 inhibition in MK led to changes in gene expression that paralleled those observed *in vivo* in platelets [39]. To assess if antiplatelet agents alter platelet expression of RUNX1 isoforms and target genes, we studied the effects of a 4-week course of daily aspirin or ticagrelor in healthy volunteers [40]. There was no effect of aspirin on platelet expression of either RUNX1 isoforms or the genes studied (Table 1). However, ticagrelor administration increased RUNX1C (logFC 0.36, p=0.037) but not RUNX1B expression (Figure 6A and Table 2). RUNX1A expression increased but did not reach statistical significance (logFC 0.38; p=0.09). *F13A1* and *PDE5A* were upregulated (Figure 6B); other genes were unchanged (Table 2). The change in RUNX1B correlated positively with the change in *F13A1* (linear estimate = 0.05, standard error = 0.02, p=0.01) and *PDE5A* (0.04, 0.03, p=0.06) (Figure 6C). Overall, these studies indicate that exposure to a P2Y12 inhibitor, but not a COX1 inhibitor, alters *in vivo* platelet expression of RUNX1 isoforms and some target genes, consistent with an effect at the MK level [39].

**Figure 6.**
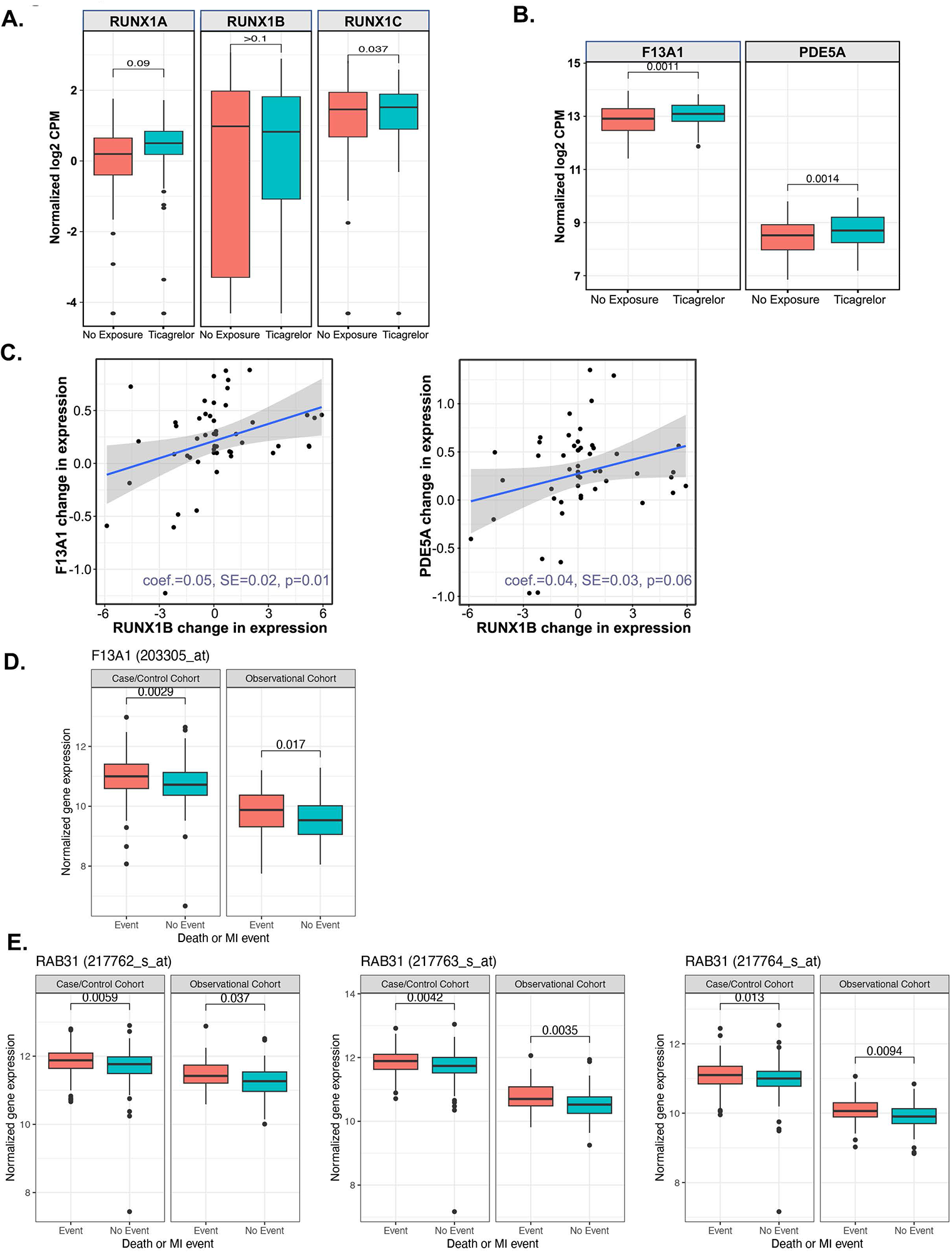
Effect of ticagrelor on expression of RUNX1 isoforms and target genes in platelets of healthy volunteers and the relationship of whole blood *F13A1* and *RAB31* levels to death or MI in patients with cardiovascular disease. **A. Effect of ticagrelor on platelet expression of RUNX1 isoforms in healthy volunteers.** Box-whisker plots of relative RUNX1 isoform gene expression quantified as TMM-normalized log2 counts per million reads mapped (log2 CPM) (x-axis) at baseline (red) and after 4 weeks of 90 mg twice daily ticagrelor (blue) exposure in healthy volunteers. Unadjusted p values generated from general linear mixed-effect modeling are presented. **B. Effects of ticagrelor exposure on platelet expression of F13A1 and PDE5A**. Box-whisker plots of relative *F13A1* and *PDE5A* gene expression were quantified as TMM-normalized log2 counts per million reads mapped (log2 CPM) (x-axis) at baseline (red) and after 4 weeks of 90 mg twice daily ticagrelor (blue) exposure in healthy volunteers. Unadjusted p values generated from general linear mixed-effect modeling are presented. **C. Correlations between changes in RUNX1B expression (x-axis) and F13A1(left panel) or PDE5A (right panel) expression (y-axis) in platelets following treatment with ticagrelor in healthy volunteers.** The correlation between RUNX1B and downstream genes *F13A1* and *PDE5A* was analyzed using linear mixed effect regression as described in the Methods section. Changes in expression are normalized log2 CPM post-treatment expression minus normalized log2 CPM pre-treatment. Points represent individual volunteers, the straight line represents the best fit linear estimate, and the shaded region represents the 95% confidence intervals. “Coef.” stands for the linear estimate; “SE” stands for the standard error. Analysis of the RNA-seq based platelet gene expression was conducted in R, using edgeR and limma for normalization and differential expression analysis, and lm or lme4 for association between gene transcripts and RUNX1 isoforms. **D. The association of *F13A1* probe set in blood with death or MI in patients with cardiovascular disease (CATHGEN cohorts).** Microarray gene expression data from two cohorts (case/control and observational cohort) using whole blood RNA collected from patients at the time of cardiac catheterization was used. Normalized expression for *F13A1* probe set is plotted in patients with death or MI outcomes (Event) vs. event free controls (No Event) in the case control and observational cohorts. The expression metrics are RMA-normalized and log transformed fluorescence intensities. Association between gene expression and the endpoint Death or MI was conducted using a moderated t-test estimated using empirical Bayes, as implemented in the R package limma. Unadjusted p values generated using empirical Bayes linear regression adjusted for age, sex, and race are presented. **E. The association of *RAB31* probe sets in blood with death or MI in patients with cardiovascular disease (CATHGEN cohorts).** Normalized expression for *RAB31* (3 probe sets used) is plotted in patients with death or MI outcomes (Event) vs. event free controls (No Event) in the case control and observational cohorts. Unadjusted p values generated using empirical Bayes linear regression adjusted for age, sex, and race are presented.

**Table 1.** Effect of Aspirin on RUNX1 isoform and target gene expression.

| **Gene** | **logFC** | **t** | **P-Value** |
| --- | --- | --- | --- |
| *RUNX1A* | -0.18 | -1.10 | 0.27 |
| *RUNX1B* | 0.44 | 1.74 | 0.08 |
| *RUNX1C* | 0.12 | 0.95 | 0.34 |
| *F13A1* | -0.01 | -0.27 | 0.79 |
| *MYL9* | -0.07 | -1.47 | 0.14 |
| *PCTP* | -0.03 | -0.31 | 0.76 |
| *PDE5A* | -0.01 | -0.08 | 0.94 |
| *RAB1B* | -0.03 | -1.15 | 0.25 |
| *RAB31* | -0.04 | -0.92 | 0.36 |

The statistical methods were described previously [39, 40]. Briefly, a linear model of gene expression (dependent variable) as a function of exposure adjusted for sex, starting RNA concentration, sequencing batch, and an adjustment for repeated measures within a given participant was estimated using the variance stabilizing weights (i.e. “voom” weights) using a moderated t-test implemented estimated using the empirical Bayes method in the R package limma. The value “t” corresponds to the moderated t-statistic. Changes in gene expression are represented as log2 fold changes. The effects are centered around 0 (i.e. 0 = no change in expression) with a symmetric interpretation, meaning an effect size of 1 corresponds to a doubling in gene expression and an effect size of -1 corresponds to a halving in gene expression. logFC, log fold change; P-Value reflect unadjusted P value.

**Table 2.**
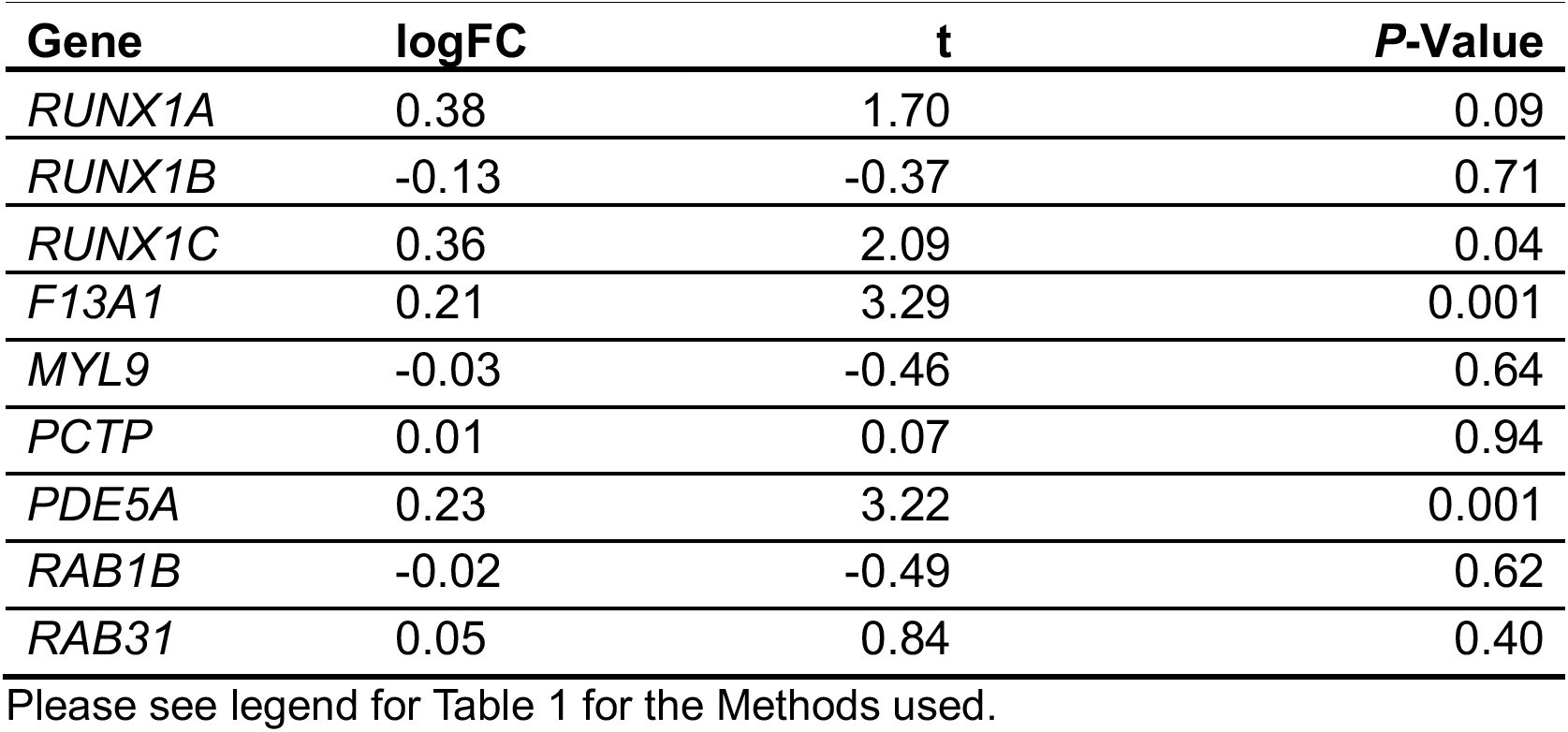
Effect of Ticagrelor on RUNX1 isoform and target gene expression.

| **Gene** | **logFC** | **t** | **P-Value** |
| --- | --- | --- | --- |
| *RUNX1A* | 0.38 | 1.70 | 0.09 |
| *RUNX1B* | -0.13 | -0.37 | 0.71 |
| *RUNX1C* | 0.36 | 2.09 | 0.04 |
| *F13A1* | 0.21 | 3.29 | 0.001 |
| *MYL9* | -0.03 | -0.46 | 0.64 |
| *PCTP* | 0.01 | 0.07 | 0.94 |
| *PDE5A* | 0.23 | 3.22 | 0.001 |
| *RAB1B* | -0.02 | -0.49 | 0.62 |
| *RAB31* | 0.05 | 0.84 | 0.40 |

Please see legend for Table 1 for the Methods used.

### Relationships to cardiovascular events

In our prior studies using Affymetrix gene arrays in patients evaluated in the cardiac catheterization laboratory and followed for 3.8 years, higher expression in whole blood of RUNX1 expressed from the P1 promoter (RUNX1C) but not the P2 promoter, was associated with protection from MI and death [29]. Higher expression of *MYL9*, *PCTP,* and *PDE5A* was associated with a higher risk of adverse cardiac events [28, 29]. We now extend these analyses to three genes - *F13A1,* and two small GTPases, *RAB31* and *RAB1B*, recently shown by us to be direct RUNX1 targets and downregulated in *RUNX1*-deficient platelets/MK [30, 33–35]. Decreased expression of *RAB31* and *RAB1B* induced defective MK vesicle trafficking (including of VWF) [34, 35]. Decreased *F13A1* expression was associated with impaired clot contraction [33]. In the present studies, higher expression of *F13A1* or *RAB31* was associated with higher CV events (Figures 6D and 6E). The effects were consistent between the two independent cohorts for both genes, and between the three probe sets available for *RAB31* (only one probe set available for *F13A1*). No such association was found for *RAB1B* (not shown).

## Discussion

Current understanding of the roles of the multiple isoforms of RUNX1 in MK and platelet biology is limited. We provide evidence that the major isoforms RUNX1B and RUNX1C autoregulate RUNX1 and regulate downstream target genes in an isoform-specific and differential manner in MK and platelets, and that this has relevance in the context of CV events, where platelets play a major role. ChIP and luciferase reporter studies (Figures 1, 2) provided evidence that RUNX1B and RUNX1C modulate P1 and P2 promoters. In HeLa cells lacking endogenous RUNX1 [15], RUNX1B inhibited and RUNX1C enhanced P1 and P2 promoter activities (Figure 2). In HEL cells, RUNX1B overexpression inhibited RUNX1A and RUNX1C expression, while RUNX1C upregulated them (Figure 2); thus, the isoforms differentially regulate RUNX1A expression also (Figure 2). Prior studies showed that RUNX1 (isoform not specified) regulated the P1 promoter [15]. We now show that both P1 and P2 promoters are regulated by RUNX1 isoforms and, of note, in an opposing manner. These findings on autoregulation have implications in attempts to therapeutically modulate endogenous RUNX1 in haplodeficient FPDMM patients [36–38, 45]. It would be important to assess the effects of therapeutic interventions on specific isoforms, including RUNX1A.

Platelets receive the bulk of transcripts from MK and reflect the MK source [46, 47]. In platelets from healthy volunteers RUNX1B transcripts by RNAseq were inversely correlated with those for RUNX1C and RUNX1A (Figure 5B); RUNX1C transcripts correlated positively with RUNX1A. There was an inverse relationship between RUNX1A and RUNX1B, both transcribed from the P2 promoter [37]. These corroborate the findings in HEL cells, providing important *in vivo* evidence of counter-regulatory effects of RUNX1 isoforms in MK.

RUNX1 regulates numerous genes in platelets and MK [28, 32–35]. In our prior studies using Affymetrix arrays, *PCTP* transcript levels in whole blood from patients were inversely related to RUNX1 transcripts from the P1 promoter [28]. We now focused on multiple genes, all direct RUNX1 targets [28–30, 32, 34, 35, 48], and provide evidence in HEL and HeLa cells that their regulation by RUNX1 isoforms is distinct with RUNX1B generally being a positive regulator (10 of 12 genes studied) and RUNX1C being a negative regulator in four and without effect on the others (Figures 3 and 4). These studies attest to the differential regulation of genes by RUNX1 isoforms.

To obtain further evidence, we explored the relationships between RUNX1 isoforms and target genes using platelet transcripts as a surrogate for MK (Figure 5). Platelet target gene expression was differentially associated with RUNX1 isoforms. RUNX1B transcripts correlated positively with six genes (*F13A1, PDE5A, PCTP, RAB1B, ALOX12,* and *PRKCQ)* and negatively with four (*MYL9, IFITM3, ITGA2B* and *PF4)*. None of these genes correlated with RUNX1C or RUNX1A. Moreover, the target genes correlated with the RUNX1B/RUNX1C ratio (Figure 5E); the direction was same as for RUNX1B. The enrichment for RUNX1B, RUNX1C, RUNX1B/RUNX1C ratio (Figure 5 F-I) and RUNX1A/RUNX1C ratio (Supplemental Figure 2B) with the downregulated genes in the RHD patient provide further corroboration.

Interestingly, in HEL cells RUNX1B was a positive regulator of *MYL9* and *ITGA2B* (Figure 4), yet in platelets, the transcripts were negatively associated (Figure 5). *PF4* may fall into this category; prior studies show that it is positively regulated by RUNX1B [49]. What may be the mechanisms? All three are robustly expressed in platelets. Studies in HEL cells reflect largely transcriptional regulation of the gene; while platelet levels reflect transcripts transferred from MK (the bulk source), and degradation and loss, including in microparticles. Not all gene transcripts are transferred to platelets to the same extent [50]. Importantly, activated platelets shed mRNA-containing, annexin-binding microparticles and constitute the predominant source of circulating microparticles [51, 52]. Disproportionate loss of *MYL9* and *ITGA2B* transcripts in microparticles relative to *RUNX1B* may yield the noted negative relationship in platelets.

Overall, the target genes are differentially regulated by RUNX1 isoforms and their expression is likely an integrated effect of RUNX1 isoforms with countervailing and competing effects. A question that arises is whether the differential regulation by RUNX1 isoforms has clinical relevance. In our previous studies [29] in patients presenting to the cardiac catheterization laboratory, higher levels in blood of transcript from the P1 promoter were protective against death or MI, and this was not noted for transcripts from the P2 promoter. These studies were performed using whole blood as platelet samples were unavailable [42]. Higher *PCTP* [28], *MYL9* [42], and *PDE5A* [42, 53] transcript levels were associated with CV events. In present studies, we examined this association for *F13A1* [33], *RAB1B* [34] and *RAB31* [35], three MK-platelet genes recently shown by us to be RUNX1 targets, downregulated in FPDMM patients and associated with abnormal platelets-MK function. *F13A1* and *RAB31* were associated with an increased incidence of death or MI in CVD patients (Figure 6). Thus, several RUNX1-target genes are associated with acute events in CVD patients.

Of these *F13A1* has important significance because it encodes for the catalytic subunit FXIII-A of coagulation factor XIII, a transglutaminase whose deficiency is associated with a bleeding disorder [54–57]. FXIII-A is synthesized by MKs (and hematopoietic cells), highly expressed in platelets (Supplemental Figure 1A) at a concentration higher than in plasma [54–56]. *F13A1* is a transcriptional target of RUNX1 [33]. Thus, although it was *F13A1* expression measured in whole blood that correlated with CV events (Figure 6D), it likely reflects an MK-platelet source. Platelet *F13A1* transcripts correlated with RUNX1B (Figure 5D). Moreover, plasma FXIII levels have been previously linked to CV events [55, 58, 59].

Our studies show distinct effects of RUNX1B and RUNX1C, which differ by only 32 AA at the N-terminus (Figure 1B); these likely confer differences in functionality [60], higher DNA binding affinity of RUNX1C [61], interactions with binding partners, and phosphorylation [62, 63]. RUNX1-isoform-specific binding co-regulators are described for RUNX1C and RUNX1A [27]. The mechanistic basis for the differential effects of the isoforms needs to be pursued. Thus, RUNX1 isoforms may regulate target gene expression by multiple mechanisms, driven by relative isoform levels, their specific effects on target genes, interactions with other binding partners, and compounded by the impact of RUNX1 autoregulation in an isoform-specific manner.

FPDMM results from heterozygous RUNX1 mutations and is associated with an increased risk of myeloid malignancies [1, 6, 10, 13, 61]; the underlying mechanisms or roles of RUNX1 isoforms in FPDMM are unclear. Most FPDMM patients have mutations in the conserved RUNT domain shared by the major isoforms [1, 64]; their relative levels are unknown in these patients, but important in understanding the alterations in platelet-MK biology and malignant transformation. Alterations in the relative expression of RUNX1 isoforms have been linked to myeloid dysplasia and malignancies. RUNX1A overexpression and altered RUNX1A/RUNX1C ratio has been linked to increased leukemia risk in trisomy 21 and Down’s syndrome [27, 65]. RUNX1A but not RUNX1B is overexpressed in CD34+ cells in myeloproliferative neoplasms [26]. Strategies to increase RUNX1 expression have been advanced as potential approaches to prevent progression to myeloid malignancies in FPDMM [36–38]. It would be relevant to assess their effects on the individual isoforms.

Exposure to ticagrelor altered *in vivo* platelet expression of RUNX1 isoforms and some target genes (Table 2), suggesting a drug-driven modulation of RUNX1. Interestingly, platelet *F13A1* and *PDE5A* transcripts increased following ticagrelor administration (Figure 6C). These likely represent a compensatory effect in response to platelet inhibition, as previously observed with respect to other genes following aspirin [40] and ticagrelor [39] administration. Moreover, studies by two of the authors (RM and DV) [39] have shown that changes in genes in CD34+-derived MK treated *in vitro* with a P2Y12 inhibitor correlated with those in platelets of healthy subjects administered ticagrelor.

Our previous studies in megakaryocytic cells *in vitro* and in whole blood of healthy individuals on a 4-week aspirin regimen showed that aspirin enhances P1 promoter-driven RUNX1 transcripts and inhibits those from the P2 promoter [29]. In the present studies, platelet RUNX1 isoforms were unaltered by aspirin (Table 1). This may be due to their low-level expression in platelets (Supplemental Figure 1A) and the relatively small effect, in general, of aspirin therapy on platelet transcripts [40].

Overall, our studies provide evidence for differential effects of RUNX1 isoforms in regulating its own expression and that of target genes in MK/platelets, and for association with acute events in CVD patients. The isoform-specific effects add to the complexity of RUNX1-driven gene regulation in MK/platelets and may be relevant to the outcomes of therapeutic approaches to alter RUNX1 expression in FPDMM.

## Acknowledgements

This study was supported by research funding from NIH (NHLBI), R01 HL137376 and R01 HL109568 to AKR.

## Author Contributions

LG performed the research, analyzed the data, and wrote the manuscript; RM analyzed the data from the CATHGEN cohorts and healthy subjects; FDC performed the research; DV conceived, designed, performed the research and interpreted data involving studies in the CATHGEN cohorts and healthy subjects; AKR conceived, designed and directed the project, participated in all aspects of the research and wrote the manuscript. All authors contributed to the manuscript, have read and approved the manuscript.

## Supplemental Materials

### Methods

#### Reagents

Dual Luciferase Reporter Assay System kit, polymerase chain reaction (PCR) reagents, Go Taq Green Master Mix, PGL4-basic vector, and the Renilla luciferase control vector were purchased from Promega Biotech (Madison, WI). RNeasy Plus Micro kit was from Qiagen (Germantown, MD). Phorbol 12-myristate 13-acetate (PMA), lipofectamine 2000 Transfection Kit, SuperScript IV First Strand Synthesis System were from Invitrogen (Waltham, MA). PowerUp SYBR Green Master Mix was from Applied Biosystems (Waltham, MA). ChIP-IT assay kit was from Active Motif (Carlsbad, CA)

#### Antibodies

Sources of antibodies: mouse-anti-RUNX1, mouse-anti-MYL9 and mouse-anti-GAPDH (Santa Cruz Biotechnology, Dallas, TX), rabbit-anti-PCTP (ThermoFisher Scientific, Waltham, MA), rabbit-anti-PDE5A (Proteintech, Rosemont, IL), sheep-anti-F13A (Affinity Biologicals, Ontario, Canada). The commercially available anti-RUNX1 antibodies recognize multiple RUNX1 isoforms. We developed a rabbit polyclonal antibody specifically against RUNX1C by targeting the 16 amino acids (MASDSIFESFPSYPQC) specific to the N-terminus of RUNX1C protein. The validation of this antibody is shown in Figure 1.

#### Cells

Human erythroleukemia (HEL) cells obtained from the American Type Culture Collection (Rockville, MD) were cultured in RPMI-1640 medium (Mediatech, Manassas, VA) supplemented with 10% FBS (GE Healthcare, Mississauga, ON, Canada) and penicillin (100 U/ml) and streptomycin (100 mg/ml) (Invitrogen, Waltham, MA) at 37°C in humidified 5% CO_2_ atmosphere. HEL cells were treated with 30 nM PMA to induce megakaryocytic transformation. Human cervical carcinoma (HeLa) cells obtained from the American Type Culture Collection (Rockville, MD), which do not express RUNX1, were grown in DMEM medium (Mediatech, Manassas, VA) supplemented with 10% FBS, penicillin and streptomycin. Cells were maintained at 37°C in humidified 5% CO_2_ atmosphere.

#### Chromatin immunoprecipitation assay

Chromatin immunoprecipitation (ChIP) assays were performed on HEL cells treated with PMA for 48 hours and then crosslinked by 1% formaldehyde for 10 minutes. ChIP analysis was performed with the Active Motif ChIP-IT assay kit. Chromatin samples were immunoprecipitated with anti-RUNX1 antibody and our anti-RUNX1C antibody and analyzed by polymerase chain reaction (PCR) using primers listed in Supplemental Table 1. Amplification was performed using GoTaq Green Master Mix and PCR products analyzed by agarose gel electrophoresis.

#### Construction of luciferase reporter plasmids

The human RUNX1 promoter regions were amplified using genomic DNA from HEL cells. The RUNX1 P1 promoter region ∼1.1kbp DNA fragment spans 596bp of 5’ upstream of exon1, 445bp of exon 1 untranslated region (UTR) and 42bp of coding sequence. The RUNX1 P2 promoter region ∼1.5kbp DNA fragment spans 1471bp of exon 3 UTR and 39bp of coding sequence. The DNA fragments were amplified using primers listed in Supplemental Table 1. They were subcloned into the PGL4-basic vector. Mutations were introduced into the five RUNX1 consensus binding sites in P1 promoter and the single site on P2 promoter using Site-Directed Mutagenesis Kit (NEB, Ipswich, MA, primers listed in Supplemental Table 1).

#### Luciferase reporter assays

HEL cells (3x10^5^) treated with 30 nM PMA for 24 hours in 12-well plates were co-transfected with vector with RUNX1-P1 or RUNX1-P2 luciferase reporter (1000 ng) and pRL-TK (20 ng) containing Renilla luciferase gene (50:1 ratio) using Lipofectamine 2000 Reagent. An empty vector, pGL4-Basic, was transfected as a control.

HeLa cells (3x10^5^) cultured in 12-well plates were co-transfected with 800 ng of P1 or P2 reporter vector, different concentrations of pCMV6-XL4-RUNX1B or M02-RUNX1C expression vector and fixed amount of pRL-TK plasmid (16 ng) using lipofectamine. After 24 hours, luciferase activity was measured in cell lysates with a Dual Luciferase Assay System. Promoter activity was expressed as firefly luciferase activity/Renilla luciferase activity relative to that of the PGL4-Basic vector.

#### Platelet RNA Quantification and Statistical Analyses

Sequence data processing and alignment is described in detail previously [1]. Briefly, Trimmomatic was used to remove adaptor and low-quality bases. Trimmed sequences were aligned to a custom ribosomal RNA database, any reads mapped to rRNA transcripts were removed and the remaining, unmapped (i.e. non-rRNA sequences) were aligned with TopHat2 v2.0.9 to the hg19 transcriptome (Ensemble v74), using both gene-level and isoform-level versions to quantify gene expression. In both cases, expression quantification was done using Cuffquant v2.2.0 and Cuffnorm v2.2.0. Cuffnorm isoform counts and gene counts were de-normalized by sample-level internal scale factors to facilitate analysis with RNA-seq count-based methods. Genes (or isoforms) with count < 10 in 80% or more samples were excluded and samples with ribosomal content > 35% of total reads or that had mean pairwise rank correlations 3 standard deviations lower than the overall mean sample-sample pairwise correlation were excluded (N = 12).

Differential expression by aspirin and ticagrelor treatment are described in detail previously [1, 2]. Briefly, gene counts and isoform counts were normalized using trimmed mean of M-values method and then variance stabilizing weights were estimated using voom, implemented in the r package *limma*. For both gene-level and isoform-level expression, expression was modeled as a function of treatment drug, controlling for sex, starting RNA concentration, flow cell, and repeated measures from individual participants. The linear model was estimated using the empirical Bayes method and generalized linear hypothesis testing (i.e. contrasts) was used to test differences between specific treatment drugs or combinations of treatment drugs.

#### Patient Cohorts with Cardiovascular Disease (CVD)

The patient cohorts with cardiovascular disease have been previously described [3, 4]. The Catheterization Genetics (CATHGEN) biorepository has banked, whole-blood RNA in PAXgene tubes from the Duke University Medical Center patients from the time of cardiac catheterization, baseline medical history, and follow-up for all-cause death and MI. CATHGEN consists of 2 cohorts of patients with CVD, an observational and a case/control cohort. In the observational cohort, 224 sequential samples were selected for RNA analysis, of which 191 had sufficient RNA for microarray analysis. The case/control cohort consisted of participants who had experienced death or MI (n=250) after their index catheterization and age-matched, sex-matched, and race-matched controls (n=250) who were free of death or MI for > 2 years after cardiac catheterization. 447 had sufficient RNA for microarray analysis; 44 overlapped with the observational cohort and were dropped, leaving 403 subjects for analysis.

Followed-up for death and MI was ascertained in both cohorts in October 2011; the median follow-up duration was 3.8 year. Patients with incomplete follow-up were censored at the time of last contact. Patients who had histories of cardiac transplantation at the time of catheterization (n=5), died within 7 days (n=1), or failed quality control (n=1) were excluded. The remaining datasets left 190 samples in the observational cohort (48 death or MI events) and 397 (202 death or MI events) in the case/control cohort.

#### Statistical analysis of data from the CATHGEN study

Statistical analysis was previously described [3]. Microarray gene expression data were normalized with Robust Multiarray Average separately for the 2 cohorts making up the CATHGEN study. To assess for the association between *F13A1* and *RAB31* with cardiovascular events, death or MI, we used longitudinal follow-up data from CATHGEN participants. In each cohort, the *F13A1* probe set (203305_at) and *RAB31* probe sets (217762_s_at, 217763_s_at and 217764_s_at) were tested for effects on MI or death events with logistic regression. Two covariate models were evaluated: the reduced covariate model controlling for the effects of age, sex, and race on death or MI outcomes and the full covariate model controlling for the effects of age, sex, race, smoking status, body mass index, hyperlipidemia, hypertension, diabetes mellitus, platelet count, aspirin treatment, and aspirin-response signature genes [5] previously shown to correlate with death or MI outcomes in these cohorts. For each probe, the log odds ratios (ORs) and their standard errors were pooled as the weighted average of the log ORs with the use of inverse variance weights, assuming a fixed-effect model the log-fold change of *F13A1* and *RAB31* expression associated with death or MI was estimated with linear regression, modeling *F13A1* and *RAB31* expression as response to death or MI outcome and adjusting for age, sex, and race. All CATHGEN data processing and statistical analyses were conducted in R version 3.2 (http://www.r-projet.org/) with limma packages for normalization, meta-analyses, and moderated t tests, respectively.

---

**Supplemental Figure 1.**
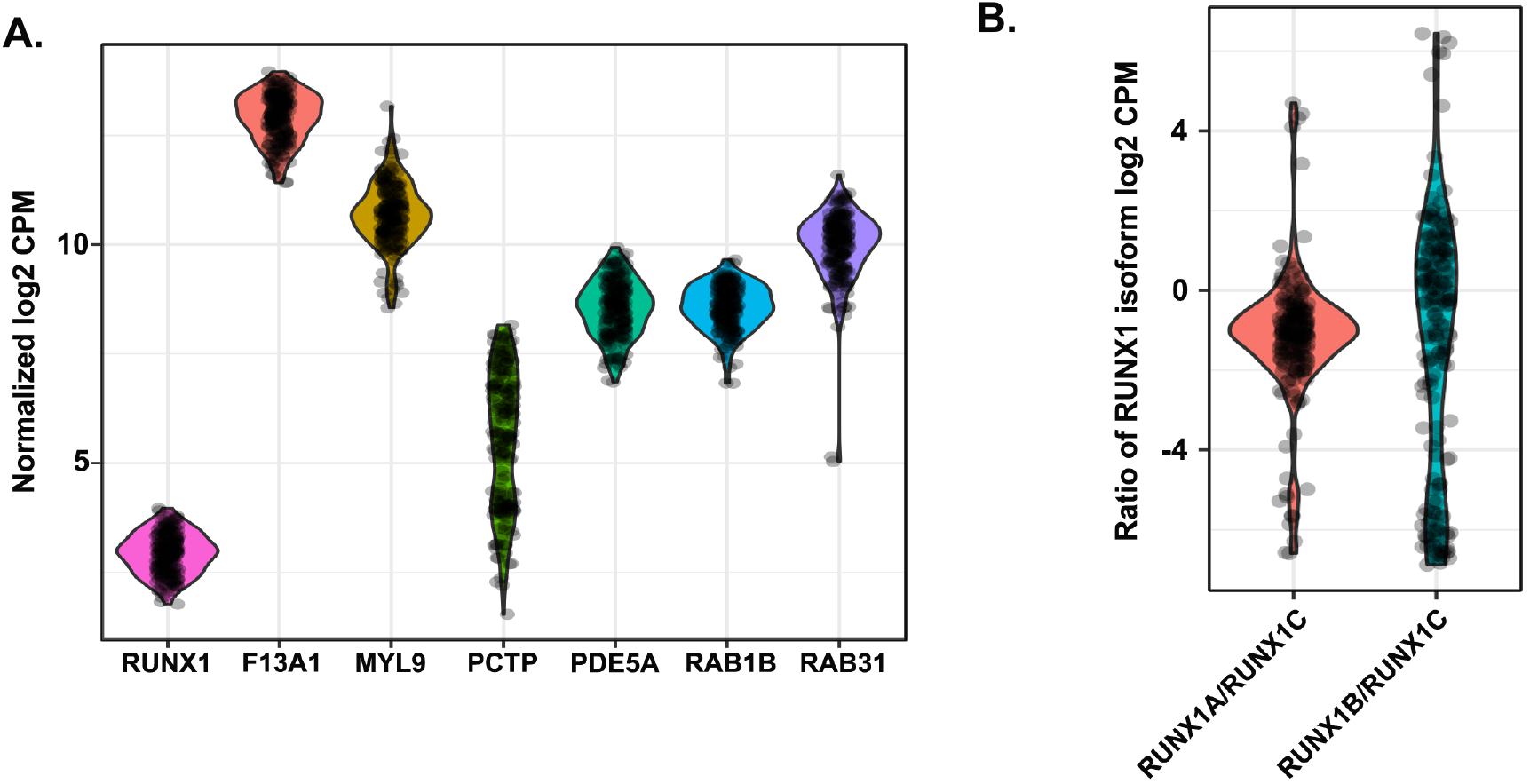
Relative expression of RUNX1 isoforms and target genes in platelets from healthy volunteers. **A. Relative expression of RUNX1 and target genes in human platelets.** RNA was isolated from leukocyte-poor platelets from 85 healthy volunteers and analyzed using RNAseq as described in the Methods. Shown are the TMM-normalized expression represented as log2 counts per million reads mapped (CPM) in baseline samples of total RUNX1 expression (i.e. all isoforms) and target genes *F13A1, MYL9, PCTP, PDE5A, RAB1B* and *RAB31*. **B. C. Ratios of RUNX1A to RUNX1C and RUNX1B to RUNX1C expression in platelets.** Ratios of the RUNX1 isoforms are represented as log2 (TMM-normalized Isoform 1 CPM/TMM-normalized isoform 2 CPM), positive ratio values indicate higher expression of isoform 1 while negative values indicate higher levels of isoform 2.

**Supplemental Figure 2.**
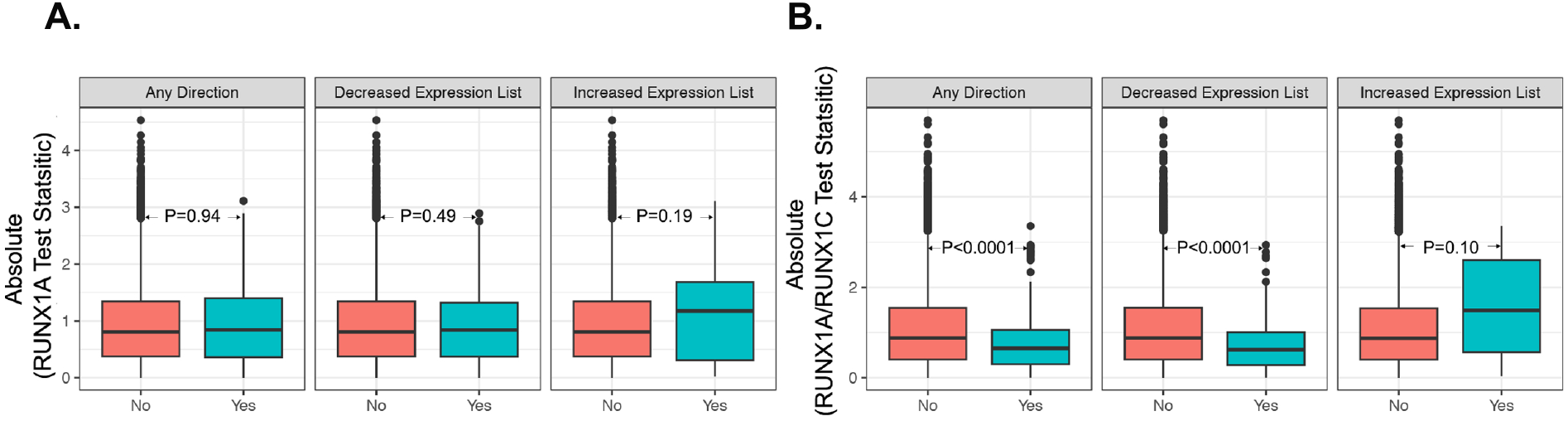
Results of Enrichment Analysis of RUNX1A and RUNX1A/RUNX1C ratio. RNA was isolated from leukocyte-poor platelets from 85 healthy volunteers and analyzed for genome wide gene expression using RNAseq. The resulting association test statistics (absolute value of the t-statistics) were used to assess the enrichment of RUNX1A (A) and RUNX1A/RUNX1C (B) correlation within RHD regulated gene sets (21 increased expression and 228 decreased expression). Shown are the absolute test statistic values (y-axis) using a two-sample t-test across three categories: Any Direction, Decreased Expression List, and Increased Expression List. “No” indicates non-RHD regulated gene set. “Yes” indicates RHD regulated gene set. Enrichment p values generated from Fisher’s exact test are presented.

**S Table 1.**
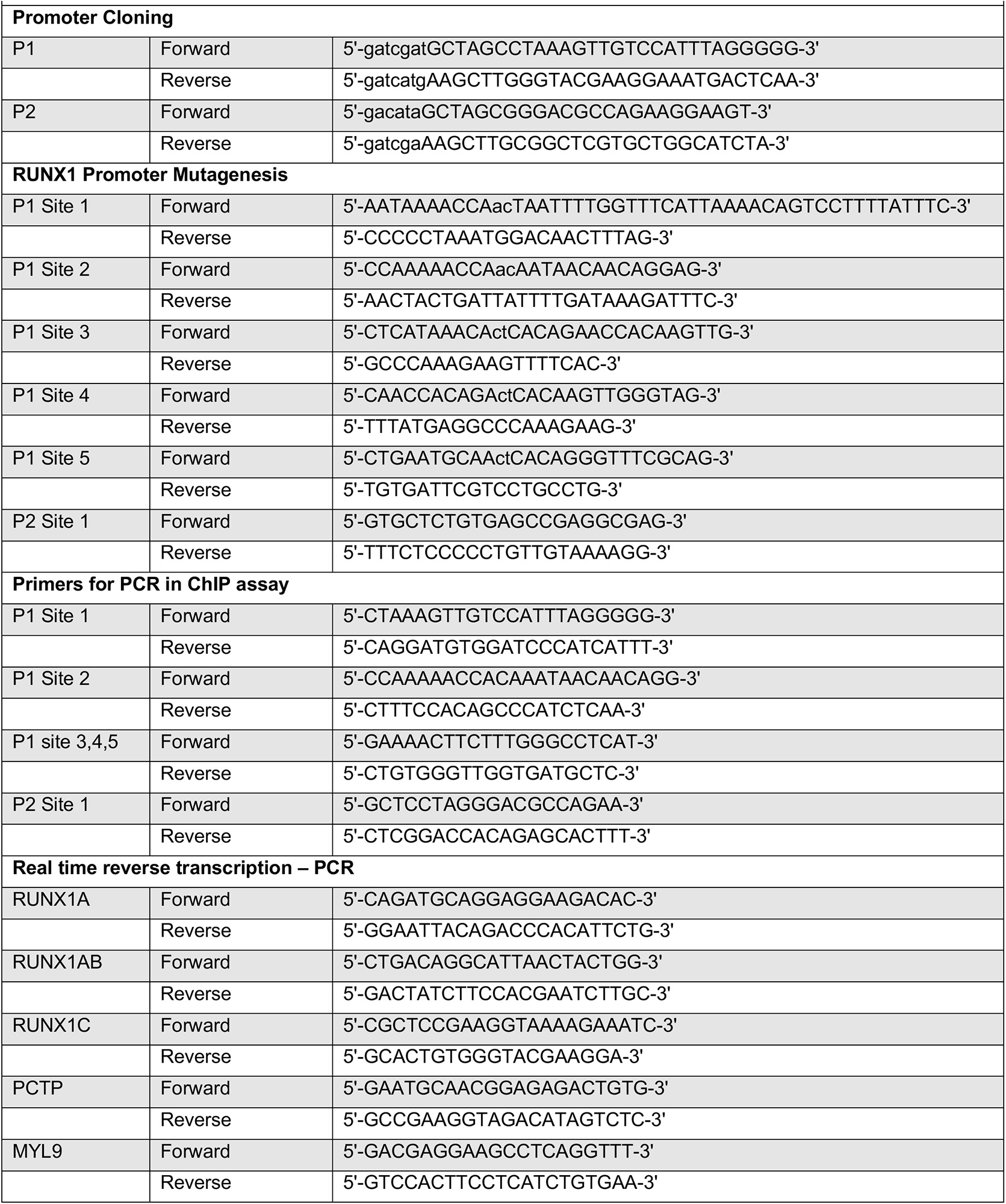
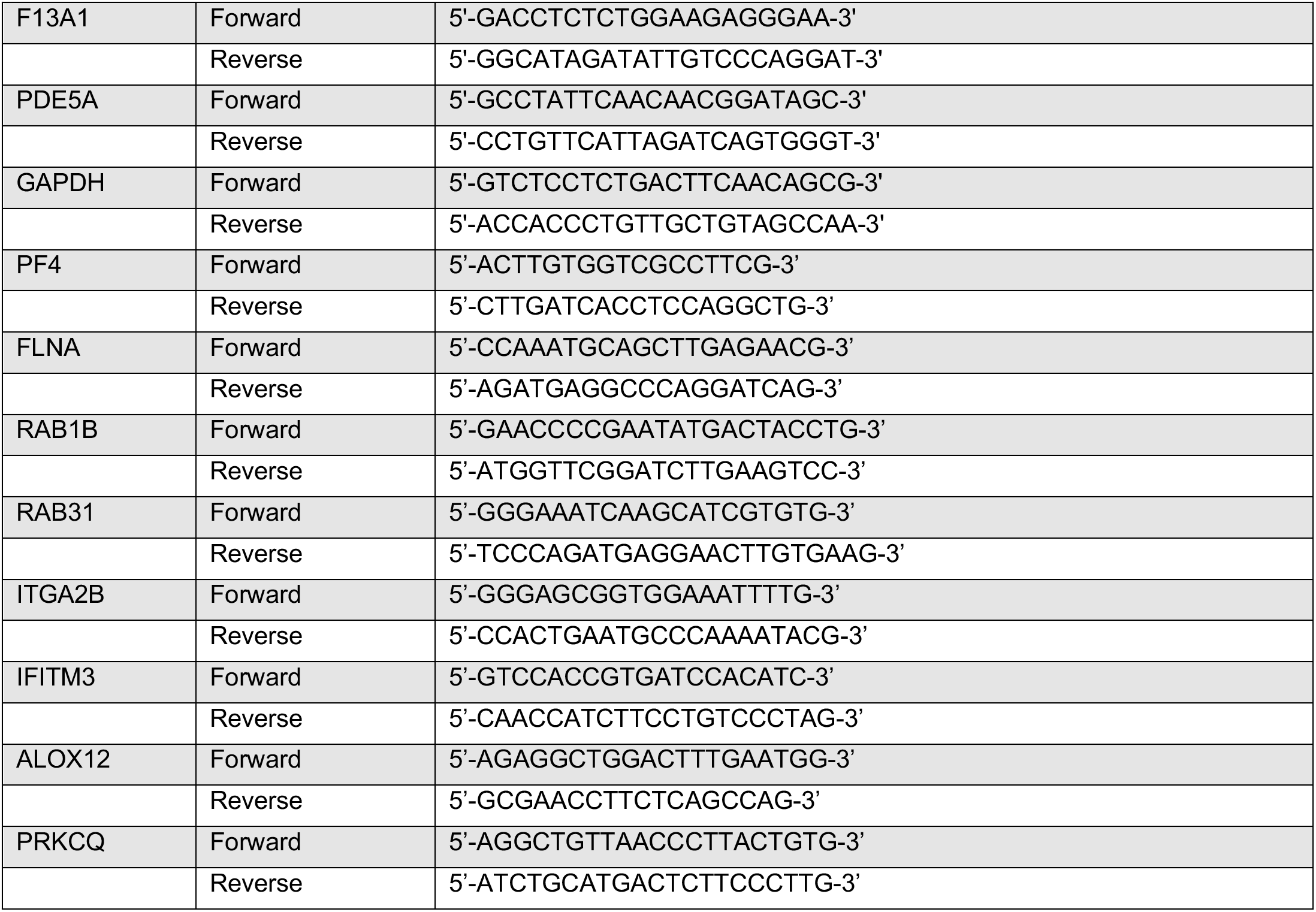
List of Primers.

**S Table 2.** Correlations between RUNX1B and Genes with Altered Expression in Platelets.

|  | Negatively correlated with RUNX1B | Positively correlated with RUNX1B |
| --- | --- | --- |
| Upregulated RHD genes (21 genes) | 3 | 0 |
| Downregulated RHD genes (228 genes) | 31 | 80 |

**S Table 3.**
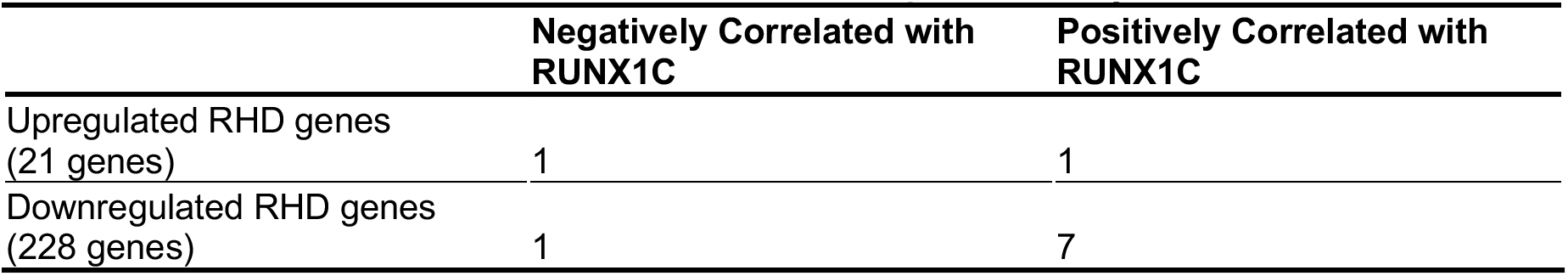
Correlations between RUNX1C and Genes with Altered Expression in Platelets from Patient with *RUNX1* Haplodeficiency.

**S Table 4.**
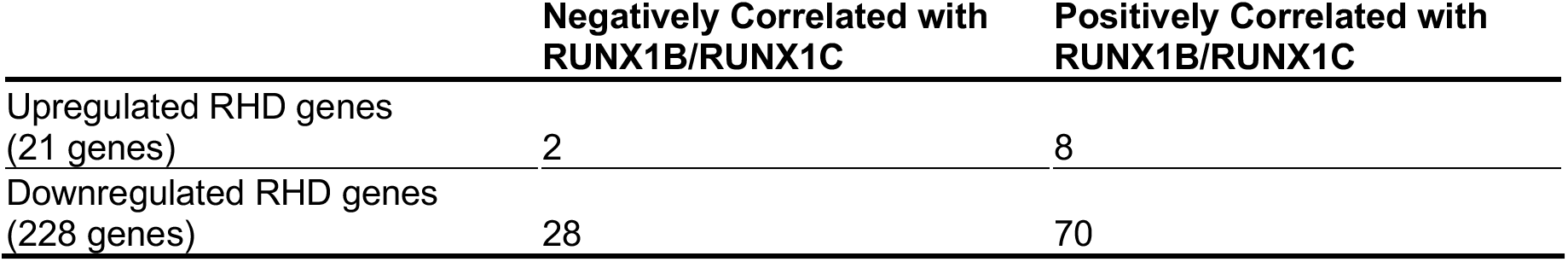
Correlation between RUN1B/RUNX1C and Genes with Altered Expression in Platelets from Patient with *RUNX1* Haplodeficiency.

## References

1 Hayashi Y, Harada Y, Harada H. Myeloid neoplasms and clonal hematopoiesis from the RUNX1 perspective. Leukemia. 2022; 36: 1203–14. 10.1038/s41375-022-01548-7.

2 de Bruijn M, Dzierzak E. Runx transcription factors in the development and function of the definitive hematopoietic system. Blood. 2017; 129: 2061–9. 10.1182/blood-2016-12-689109.

3 Ichikawa M, Asai T, Chiba S, Kurokawa M, Ogawa S. Runx1/AML-1 ranks as a master regulator of adult hematopoiesis. Cell Cycle. 2004; 3: 722–4.

4 Liau WS, Ngoc PC, Sanda T. Roles of the RUNX1 Enhancer in Normal Hematopoiesis and Leukemogenesis. Adv Exp Med Biol. 2017; 962: 139–47. 10.1007/978-981-10-3233-2_10.

5 Tracey WD, Speck NA. Potential roles for RUNX1 and its orthologs in determining hematopoietic cell fate. Semin Cell Dev Biol. 2000; 11: 337–42. 10.1006/scdb.2000.0186.

6 Cunningham L, Merguerian M, Calvo KR, Davis J, Deuitch NT, Dulau-Florea A, Patel N, Yu K, Sacco K, Bhattacharya S, Passi M, Ozkaya N, De Leon S, Chong S, Craft K, Diemer J, Bresciani E, O’Brien K, Andrews EJ, Park N, Hathaway L, Cowen EW, Heller T, Ryan K, Barochia A, Nghiem K, Niemela J, Rosenzweig S, Young DJ, Frischmeyer-Guerrerio PA, Braylan R, Liu PP. Natural history study of patients with familial platelet disorder with associated myeloid malignancy. Blood. 2023; 142: 2146–58. 10.1182/blood.2023019746.

7 Sood R, Kamikubo Y, Liu P. Role of RUNX1 in hematological malignancies. Blood. 2017; 129: 2070–82. 10.1182/blood-2016-10-687830.

8 Song WJ, Sullivan MG, Legare RD, Hutchings S, Tan X, Kufrin D, Ratajczak J, Resende IC, Haworth C, Hock R, Loh M, Felix C, Roy DC, Busque L, Kurnit D, Willman C, Gewirtz AM, Speck NA, Bushweller JH, Li FP, Gardiner K, Poncz M, Maris JM, Gilliland DG. Haploinsufficiency of CBFA2 causes familial thrombocytopenia with propensity to develop acute myelogenous leukaemia. Nat Genet. 1999; 23: 166–75. 10.1038/13793.

9 Songdej N, Rao AK. Hematopoietic transcription factor mutations: important players in inherited platelet defects. Blood. 2017; 129: 2873–81. 10.1182/blood-2016-11-709881.

10 Deuitch N, Broadbridge E, Cunningham L, Liu P. RUNX1 Familial Platelet Disorder with Associated Myeloid Malignancies. In: Adam MP, Feldman J, Mirzaa GM, Pagon RA, Wallace SE, Bean LJH, Gripp KW, Amemiya A, eds. GeneReviews((R)). Seattle (WA), 1993.

11 Ghozi MC, Bernstein Y, Negreanu V, Levanon D, Groner Y. Expression of the human acute myeloid leukemia gene AML1 is regulated by two promoter regions. Proc Natl Acad Sci U S A. 1996; 93: 1935–40. 10.1073/pnas.93.5.1935.

12 Ran D, Shia WJ, Lo MC, Fan JB, Knorr DA, Ferrell PI, Ye Z, Yan M, Cheng L, Kaufman DS, Zhang DE. RUNX1a enhances hematopoietic lineage commitment from human embryonic stem cells and inducible pluripotent stem cells. Blood. 2013; 121: 2882–90. 10.1182/blood-2012-08-451641.

13 Lie ALM, Mevel R, Patel R, Blyth K, Baena E, Kouskoff V, Lacaud G. RUNX1 Dosage in Development and Cancer. Mol Cells. 2020; 43: 126–38. 10.14348/molcells.2019.0301.

14 Draper JE, Sroczynska P, Tsoulaki O, Leong HS, Fadlullah MZ, Miller C, Kouskoff V, Lacaud G. Correction: RUNX1B Expression Is Highly Heterogeneous and Distinguishes Megakaryocytic and Erythroid Lineage Fate in Adult Mouse Hematopoiesis. PLoS Genet. 2016; 12: e1006084. 10.1371/journal.pgen.1006084.

15 Martinez M, Hinojosa M, Trombly D, Morin V, Stein J, Stein G, Javed A, Gutierrez SE. Transcriptional Auto-Regulation of RUNX1 P1 Promoter. PloS one. 2016; 11: e0149119. 10.1371/journal.pone.0149119.

16 Swiers G, de Bruijn M, Speck NA. Hematopoietic stem cell emergence in the conceptus and the role of Runx1. Int J Dev Biol. 2010; 54: 1151–63. 10.1387/ijdb.103106gs.

17 Mikhail FM, Sinha KK, Saunthararajah Y, Nucifora G. Normal and transforming functions of RUNX1: a perspective. J Cell Physiol. 2006; 207: 582–93.

18 Leong WY, Guo H, Ma O, Huang H, Cantor AB, Friedman AD. Runx1 Phosphorylation by Src Increases Trans-activation via Augmented Stability, Reduced Histone Deacetylase (HDAC) Binding, and Increased DNA Affinity, and Activated Runx1 Favors Granulopoiesis. J Biol Chem. 2016; 291: 826–36. 10.1074/jbc.M115.674234.

19 Boisset JC, van Cappellen W, Andrieu-Soler C, Galjart N, Dzierzak E, Robin C. In vivo imaging of haematopoietic cells emerging from the mouse aortic endothelium. Nature. 2010; 464: 116–20. 10.1038/nature08764.

20 Lie ALM, Marinopoulou E, Lilly AJ, Challinor M, Patel R, Lancrin C, Kouskoff V, Lacaud G. Regulation of RUNX1 dosage is crucial for efficient blood formation from hemogenic endothelium. Development. 2018; 145. 10.1242/dev.149419.

21 Challen GA, Goodell MA. Runx1 isoforms show differential expression patterns during hematopoietic development but have similar functional effects in adult hematopoietic stem cells. Exp Hematol. 2010; 38: 403–16. 10.1016/j.exphem.2010.02.011.

22 Menegatti S, Potts B, Garcia-Alegria E, Paredes R, Lie ALM, Lacaud G, Kouskoff V. The RUNX1b Isoform Defines Hemogenic Competency in Developing Human Endothelial Cells. Front Cell Dev Biol. 2021; 9: 812639. 10.3389/fcell.2021.812639.

23 Navarro-Montero O, Ayllon V, Lamolda M, Lopez-Onieva L, Montes R, Bueno C, Ng E, Guerrero-Carreno X, Romero T, Romero-Moya D, Stanley E, Elefanty A, Ramos-Mejia V, Menendez P, Real PJ. RUNX1c Regulates Hematopoietic Differentiation of Human Pluripotent Stem Cells Possibly in Cooperation with Proinflammatory Signaling. Stem Cells. 2017; 35: 2253–66. 10.1002/stem.2700.

24 Draper JE, Sroczynska P, Leong HS, Fadlullah MZH, Miller C, Kouskoff V, Lacaud G. Mouse RUNX1C regulates premegakaryocytic/erythroid output and maintains survival of megakaryocyte progenitors. Blood. 2017; 130: 271–84. 10.1182/blood-2016-06-723635.

25 Ran D, Lam K, Shia WJ, Lo MC, Fan JB, Knorr DA, Ferrell PI, Ye Z, Yan M, Cheng L, Kaufman DS, Zhang DE. Response: the role of RUNX1 isoforms in hematopoietic commitment of human pluripotent stem cells. Blood. 2013; 121: 5252–3. 10.1182/blood-2013-04-494914.

26 Sakurai H, Harada Y, Ogata Y, Kagiyama Y, Shingai N, Doki N, Ohashi K, Kitamura T, Komatsu N, Harada H. Overexpression of RUNX1 short isoform has an important role in the development of myelodysplastic/myeloproliferative neoplasms. Blood Adv. 2017; 1: 1382–6. 10.1182/bloodadvances.2016002725.

27 Gialesaki S, Brauer-Hartmann D, Issa H, Bhayadia R, Alejo-Valle O, Verboon L, Schmell AL, Laszig S, Regenyi E, Schuschel K, Labuhn M, Ng M, Winkler R, Ihling C, Sinz A, Glass M, Huttelmaier S, Matzk S, Schmid L, Struwe FJ, Kadel SK, Reinhardt D, Yaspo ML, Heckl D, Klusmann JH. RUNX1 isoform disequilibrium promotes the development of trisomy 21-associated myeloid leukemia. Blood. 2023; 141: 1105–18. 10.1182/blood.2022017619.

28 Mao G, Songdej N, Voora D, Goldfinger LE, Del Carpio-Cano FE, Myers RA, Rao AK. Transcription Factor RUNX1 Regulates Platelet PCTP (Phosphatidylcholine Transfer Protein): Implications for Cardiovascular Events: Differential Effects of RUNX1 Variants. Circulation. 2017; 136: 927–39. 10.1161/CIRCULATIONAHA.116.023711.

29 Voora D, Rao AK, Jalagadugula GS, Myers R, Harris E, Ortel TL, Ginsburg GS. Systems Pharmacogenomics Finds RUNX1 Is an Aspirin-Responsive Transcription Factor Linked to Cardiovascular Disease and Colon Cancer. EBioMedicine. 2016; 11: 157–64. 10.1016/j.ebiom.2016.08.021.

30 Sun L, Gorospe JR, Hoffman EP, Rao AK. Decreased platelet expression of myosin regulatory light chain polypeptide (MYL9) and other genes with platelet dysfunction and CBFA2/RUNX1 mutation: insights from platelet expression profiling. J Thromb Haemost. 2007; 5: 146–54.

31 Palma-Barqueros V, Bastida JM, Lopez Andreo MJ, Zamora-Canovas A, Zaninetti C, Ruiz-Pividal JF, Bohdan N, Padilla J, Teruel-Montoya R, Marin-Quilez A, Revilla N, Sanchez-Fuentes A, Rodriguez-Alen A, Benito R, Vicente V, Iturbe T, Greinacher A, Lozano ML, Rivera J, Grupo Espanol de Alteraciones Plaquetarias C, Spanish Society of T, Haemostasis. Platelet transcriptome analysis in patients with germline RUNX1 mutations. J Thromb Haemost. 2023; 21: 1352–65. 10.1016/j.jtha.2023.01.023.

32 Jalagadugula G, Mao G, Kaur G, Goldfinger LE, Dhanasekaran DN, Rao AK. Regulation of platelet myosin light chain (*MYL9*) by RUNX1: implications for thrombocytopenia and platelet dysfunction in *RUNX1* haplodeficiency. Blood. 2010; 116: 6037–45.

33 Del Carpio-Cano F, Songdej N, Mao G, Wurtzel J, Goldfinger L, Lambert M, Rao A. Transcription factor RUNX1 regulates Factor XIIIA subunit (F13A1) expression in platelets and megakaryocytic cells: decreased platelet F13A1 expression and clot retraction in RUNX1 haplodeficiency [abstract]. Res Pract Thromb Haemost. 2021; 2021;Supplement 5:OC 31.1.

34 Jalagadugula G, Goldfinger LE, Mao G, Lambert MP, Rao AK. Defective RAB1B-related megakaryocytic ER-to-Golgi transport in RUNX1 haplodeficiency: impact on von Willebrand factor. Blood Adv. 2018; 2: 797–806. 10.1182/bloodadvances.2017014274.

35 Jalagadugula G, Mao G, Goldfinger LE, Wurtzel J, Del Carpio-Cano F, Lambert MP, Estevez B, French DL, Poncz M, Rao AK. Defective RAB31-mediated megakaryocytic early endosomal trafficking of VWF, EGFR, and M6PR in RUNX1 deficiency. Blood Adv. 2022; 6: 5100–12. 10.1182/bloodadvances.2021006945.

36 Krutein MC, Hart MR, Anderson DJ, Jeffery J, Kotini AG, Dai J, Chien S, DelPriore M, Borst S, Maguire JA, French DL, Gadue P, Papapetrou EP, Keel SB, Becker PS, Horwitz MS. Restoring RUNX1 deficiency in RUNX1 familial platelet disorder by inhibiting its degradation. Blood Adv. 2021; 5: 687–99. 10.1182/bloodadvances.2020002709.

37 Lee K, Ahn HS, Estevez B, Poncz M. RUNX1-deficient human megakaryocytes demonstrate thrombopoietic and platelet half-life and functional defects. Blood. 2023; 141: 260–70. 10.1182/blood.2022017561.

38 Estevez B, Borst S, Jarocha D, Sudunagunta V, Gonzalez M, Garifallou J, Hakonarson H, Gao P, Tan K, Liu P, Bagga S, Holdreith N, Tong W, Speck N, French DL, Gadue P, Poncz M. RUNX-1 haploinsufficiency causes a marked deficiency of megakaryocyte-biased hematopoietic progenitor cells. Blood. 2021; 137: 2662–75. 10.1182/blood.2020006389.

39 Myers RA, Ortel TL, Waldrop A, Cornwell M, Newman JD, Levy NK, Barrett TJ, Ruggles K, Sowa MA, Dave S, Ginsburg GS, Berger JS, Voora D. Platelet RNA Biomarker of Ticagrelor-Responsive Genes Is Associated With Platelet Function and Cardiovascular Events. Arterioscler Thromb Vasc Biol. 2024; 44: 423–34. 10.1161/ATVBAHA.123.319759.

40 Myers RA, Ortel TL, Waldrop A, Dave S, Ginsburg GS, Voora D. Aspirin effects on platelet gene expression are associated with a paradoxical, increase in platelet function. Br J Clin Pharmacol. 2022; 88: 2074–83. 10.1111/bcp.15127.

41 Kondkar AA, Bray MS, Leal SM, Nagalla S, Liu DJ, Jin Y, Dong JF, Ren Q, Whiteheart SW, Shaw C, Bray PF. VAMP8/endobrevin is overexpressed in hyperreactive human platelets: suggested role for platelet microRNA. J Thromb Haemost. 2010; 8: 369–78.

42 Voora D, Cyr D, Lucas J, Chi JT, Dungan J, McCaffrey TA, Katz R, Newby LK, Kraus WE, Becker RC, Ortel TL, Ginsburg GS. Aspirin exposure reveals novel genes associated with platelet function and cardiovascular events. J Am Coll Cardiol. 2013; 62: 1267–76. 10.1016/j.jacc.2013.05.073.

43 Tijssen MR, Cvejic A, Joshi A, Hannah RL, Ferreira R, Forrai A, Bellissimo DC, Oram SH, Smethurst PA, Wilson NK, Wang X, Ottersbach K, Stemple DL, Green AR, Ouwehand WH, Gottgens B. Genome-wide analysis of simultaneous GATA1/2, RUNX1, FLI1, and SCL binding in megakaryocytes identifies hematopoietic regulators. Developmental cell. 2011; 20: 597–609. 10.1016/j.devcel.2011.04.008.

44 Zauli G, Bassini A, Catani L, Gibellini D, Celeghini C, Borgatti P, Caramelli E, Guidotti L, Capitani S. PMA-induced megakaryocytic differentiation of HEL cells is accompanied by striking modifications of protein kinase C catalytic activity and isoform composition at the nuclear level. Br J Haematol. 1996; 92: 530–6.

45 Horwitz M. Hypermethylated myoblasts specifically deficient in MyoD autoactivation as a consequence of instability of MyoD. Exp Cell Res. 1996; 226: 170–82. 10.1006/excr.1996.0216.

46 Davizon-Castillo P, Rowley JW, Rondina MT. Megakaryocyte and Platelet Transcriptomics for Discoveries in Human Health and Disease. Arterioscler Thromb Vasc Biol. 2020; 40: 1432–40. 10.1161/ATVBAHA.119.313280.

47 Esparza O, Higa K, Davizon-Castillo P. Molecular and functional characteristics of megakaryocytes and platelets in aging. Curr Opin Hematol. 2020; 27: 302–10. 10.1097/MOH.0000000000000601.

48 De Arcangelis V, De Angelis L, Barbagallo F, Campolo F, de Oliveira do Rego AG, Pellegrini M, Naro F, Giorgi M, Monaco L. Phosphodiesterase 5a Signalling in Skeletal Muscle Pathophysiology. Int J Mol Sci. 2022; 24. 10.3390/ijms24010703.

49 Aneja K, Jalagadugula G, Mao G, Singh A, Rao AK. Mechanism of platelet factor 4 (PF4) deficiency with RUNX1 haplodeficiency: RUNX1 is a transcriptional regulator of PF4. J Thromb Haemost. 2011; 9: 383–91. 10.1111/j.1538-7836.2010.04154.x.

50 Cecchetti L, Tolley ND, Michetti N, Bury L, Weyrich AS, Gresele P. Megakaryocytes differentially sort mRNAs for matrix metalloproteinases and their inhibitors into platelets: a mechanism for regulating synthetic events. Blood. 2011; 118: 1903–11. 10.1182/blood-2010-12-324517.

51 Boilard E, Bellio M. Platelet extracellular vesicles and the secretory interactome join forces in health and disease. Immunol Rev. 2022; 312: 38–51. 10.1111/imr.13119.

52 Sartori MT, Zurlo C, Bon M, Bertomoro A, Bendo R, Bertozzi I, Radu CM, Campello E, Simioni P, Fabris F. Platelet-Derived Microparticles Bearing PF4 and Anti-GAGS Immunoglobulins in Patients with Sepsis. Diagnostics (Basel). 2020; 10. 10.3390/diagnostics10090627.

53 Voora D, Coles A, Lee KL, Hoffmann U, Wingrove JA, Rhees B, Huang L, Daniels SE, Monane M, Rosenberg S, Shah SH, Kraus WE, Ginsburg GS, Douglas PS. An age- and sex-specific gene expression score is associated with revascularization and coronary artery disease: Insights from the Prospective Multicenter Imaging Study for Evaluation of Chest Pain (PROMISE) trial. Am Heart J. 2017; 184: 133–40. 10.1016/j.ahj.2016.11.004.

54 Mitchell JL, Mutch NJ. Novel aspects of platelet factor XIII function. Thromb Res. 2016; 141 Suppl 2: S17–21. 10.1016/S0049-3848(16)30356-5.

55 Wolberg AS, Sang Y. Fibrinogen and Factor XIII in Venous Thrombosis and Thrombus Stability. Arterioscler Thromb Vasc Biol. 2022; 42: 931–41. 10.1161/ATVBAHA.122.317164.

56 Alshehri FSM, Whyte CS, Mutch NJ. Factor XIII-A: An Indispensable “Factor” in Haemostasis and Wound Healing. Int J Mol Sci. 2021; 22. 10.3390/ijms22063055.

57 Muszbek L, Bereczky Z, Bagoly Z, Komaromi I, Katona E. Factor XIII: a coagulation factor with multiple plasmatic and cellular functions. Physiol Rev. 2011; 91: 931–72. 10.1152/physrev.00016.2010.

58 Bereczky Z, Balogh E, Katona E, Czuriga I, Edes I, Muszbek L. Elevated factor XIII level and the risk of myocardial infarction in women. Haematologica. 2007; 92: 287–8. 10.3324/haematol.10647.

59 Muszbek L, Bereczky Z, Bagoly Z, Shemirani AH, Katona E. Factor XIII and atherothrombotic diseases. Semin Thromb Hemost. 2010; 36: 18–33. 10.1055/s-0030-1248721.

60 Nieke S, Yasmin N, Kakugawa K, Yokomizo T, Muroi S, Taniuchi I. Unique N-terminal sequences in two Runx1 isoforms are dispensable for Runx1 function. BMC Dev Biol. 2017; 17: 14. 10.1186/s12861-017-0156-y.

61 Telfer JC, Rothenberg EV. Expression and function of a stem cell promoter for the murine CBFalpha2 gene: distinct roles and regulation in natural killer and T cell development. Dev Biol. 2001; 229: 363–82. 10.1006/dbio.2000.9991.

62 Huang H, Woo AJ, Waldon Z, Schindler Y, Moran TB, Zhu HH, Feng GS, Steen H, Cantor AB. A Src family kinase-Shp2 axis controls RUNX1 activity in megakaryocyte and T-lymphocyte differentiation. Genes Dev. 2012; 26: 1587–601. 10.1101/gad.192054.112.

63 Neel BG, Speck NA. Tyrosyl phosphorylation toggles a Runx1 switch. Genes Dev. 2012; 26: 1520–6. 10.1101/gad.198051.112.

64 Yu K, Deuitch N, Merguerian M, Cunningham L, Davis J, Bresciani E, Diemer J, Andrews E, Young A, Donovan F, Sood R, Craft K, Chong S, Chandrasekharappa S, Mullikin J, Liu PP. Genomic Landscape of Patients with Germline RUNX1 Variants and Familial Platelet Disorder with Myeloid Malignancy. bioRxiv. 2023. 10.1101/2023.01.17.524290.

65 Rozen EJ, Ozeroff CD, Allen MA. RUN(X) out of blood: emerging RUNX1 functions beyond hematopoiesis and links to Down syndrome. Hum Genomics. 2023; 17: 83. 10.1186/s40246-023-00531-2.

## References

1 Myers RA, Ortel TL, Waldrop A, Dave S, Ginsburg GS, Voora D. Aspirin effects on platelet gene expression are associated with a paradoxical, increase in platelet function. Br J Clin Pharmacol. 2022; 88: 2074–83. 10.1111/bcp.15127.

2 Myers RA, Ortel TL, Waldrop A, Cornwell M, Newman JD, Levy NK, Barrett TJ, Ruggles K, Sowa MA, Dave S, Ginsburg GS, Berger JS, Voora D. Platelet RNA Biomarker of Ticagrelor-Responsive Genes Is Associated With Platelet Function and Cardiovascular Events. Arterioscler Thromb Vasc Biol. 2024; 44: 423–34. 10.1161/ATVBAHA.123.319759.

3 Mao G, Songdej N, Voora D, Goldfinger LE, Del Carpio-Cano FE, Myers RA, Rao AK. Transcription Factor RUNX1 Regulates Platelet PCTP (Phosphatidylcholine Transfer Protein): Implications for Cardiovascular Events: Differential Effects of RUNX1 Variants. Circulation. 2017; 136: 927–39. 10.1161/CIRCULATIONAHA.116.023711.

4 Voora D, Cyr D, Lucas J, Chi JT, Dungan J, McCaffrey TA, Katz R, Newby LK, Kraus WE, Becker RC, Ortel TL, Ginsburg GS. Aspirin exposure reveals novel genes associated with platelet function and cardiovascular events. J Am Coll Cardiol. 2013; 62: 1267–76. 10.1016/j.jacc.2013.05.073.

5 Voora D, Rao AK, Jalagadugula GS, Myers R, Harris E, Ortel TL, Ginsburg GS. Systems Pharmacogenomics Finds RUNX1 Is an Aspirin-Responsive Transcription Factor Linked to Cardiovascular Disease and Colon Cancer. EBioMedicine. 2016; 11: 157–64. 10.1016/j.ebiom.2016.08.021.

